# Substrate identification and specificity profiling of deubiquitylases against endogenously-generated ubiquitin-protein conjugates

**DOI:** 10.1101/2023.12.20.572581

**Authors:** Valentina Rossio, Joao A. Paulo, Xinyue Liu, Steven P. Gygi, Randall W. King

## Abstract

Deubiquitylating enzymes (DUBs) remove ubiquitin from proteins thereby regulating their stability or activity. Our understanding of DUB-substrate specificity is limited because DUBs are typically not compared to each other against many physiological substrates. By broadly inhibiting DUBs in *Xenopus* egg extract, we generated hundreds of ubiquitylated proteins and compared the ability of 30 DUBs to deubiquitylate them using quantitative proteomics. We identified five high impact DUBs (USP7, USP9X, USP36, USP15 and USP24) that each reduced ubiquitylation of over ten percent of the isolated proteins. Candidate substrates of high impact DUBs showed substantial overlap and were enriched for disordered regions, suggesting this feature may promote substrate recognition. Other DUBs showed lower impact and non-overlapping specificity, targeting distinct non-disordered proteins including complexes such as the ribosome or the proteasome. Altogether our study identifies candidate DUB substrates and defines patterns of functional redundancy and specificity, revealing substrate characteristics that may influence DUB-substrate recognition.

## Introduction

Ubiquitylation is a versatile and reversible post-translational modification that controls many cellular processes^1–4^. Installed by a cascade of enzymes (E1, E2, and E3), ubiquitin can be attached to substrates as single molecules or as ubiquitin chains of different topology to confer specific functions^1–4^. The best characterized function of ubiquitylation is to target proteins for degradation by the proteasome^3,5^. However, ubiquitylation can also alter the activity or location of proteins without targeting them for degradation, regulating many cellular pathways including protein trafficking, DNA repair and ribosomal function^6^.

Conversely, deubiquitylating enzymes (DUBs) remove ubiquitin from proteins, controlling their stability or activity. DUBs are therefore essential for suppressing proteasomal degradation of certain substrates and for modulating the functions of proteins regulated by non-proteolytic ubiquitylation. Furthermore, DUBs maintain availability of free ubiquitin to ensure the activity of the ubiquitin-proteasome system. There are roughly 100 DUBs encoded in the human genome, classified into two main categories based on their mechanism of action: a large family of cysteine-proteases and a small family of zinc-dependent metalloproteases. The cysteine-protease DUBs are further subdivided into several families including USP, OTU, MINDY, MJD, and ZUFSP. The USP family is the largest, containing 56 DUBs^7–9^.

A major challenge in the field is to identify substrates for each DUB, which is necessary to understand the mechanisms that explain substrate selectivity and for characterizing the cellular roles of these enzymes. The number of substrates identified for each DUB is heterogeneous: some well-studied DUBs have hundreds of known substrates whereas some poorly studied enzymes have none^10^. Identifying DUB substrates is challenging because inactivation of a single DUB may not be sufficient to alter stability or ubiquitylation of a substrate due to functional redundancy^11–13^. In contrast, some DUBs are essential, complicating the performance and interpretation of loss-of-function experiments. Because DUBs are often characterized individually in different experimental systems, we have little understanding of the relative physiological specificity of these enzymes.

Most of the knowledge about DUB selectivity has been acquired by monitoring the activity of recombinant DUBs on model substrates consisting of di-ubiquitin or ubiquitin chains of defined topology, or ubiquitin attached to different leaving groups^14–20^. These assays revealed that some enzymes, such as the OTU DUB family, bind ubiquitin chains of specific topologies and hydrolyze specific ubiquitin-ubiquitin linkages^19^. Other enzymes, in particular the USP DUB family, cleave ubiquitin from the substrate in a non-chain-selective manner by binding substrates directly^7,9,16^. Therefore, developing methods for identifying the substrate proteins to which ubiquitin is attached and removed, and how DUBs influence this process, is critical for understanding the principles that underlie the functional selectivity of DUBs.

In previous research^13^, we overcame the challenge posed by redundant action of DUBs by broadly inhibiting cysteine-protease DUBs simultaneously using ubiquitin vinyl-sulfone (UbVS)^21^. This led to the discovery of a set of proteins whose stability relies on DUB activity. Here, using a related strategy, we generated hundreds of proteins modified by non-proteolytic ubiquitylation. By comparing the activity of thirty DUBs against these proteins, we uncovered distinct patterns of activity that highlight both specific as well as redundant action of these enzymes. We found that a small set of highly active DUBs act in a partially redundant fashion on a wide range of substrates, whereas a distinct set of DUBs act on smaller, separate pools of substrates. This analysis identified candidate substrates for many DUBs, as well as ubiquitylated proteins that were insensitive to all tested DUBs. Candidate substrates were enriched in disordered regions compared to proteins that were insensitive to DUBs, suggesting that this feature might promote DUB recognition and in particular recognition by high impact DUBs. Furthermore, our study identified many DUB substrates previously reported in human cells, suggesting that our method can reveal *bona fide* substrates for DUBs that are regulated by ubiquitylation in different experimental systems.

## Results

### Impact and Effect of DUBs on the ubiquitylated proteome

We established a quantitative proteomic approach to compare DUB activity across hundreds of ubiquitylated proteins in a cell lysate. We treated *Xenopus* egg extract with UbVS^21^ and after UbVS was consumed by reaction with endogenous DUBs, we added back single recombinant enzymes, together with HA-tagged ubiquitin (HA-Ub), which is incorporated into HA-Ub-conjugates (Fig.1A). We then measured with TMT-based quantitative proteomics how the abundance of the immunopurified HA-tagged species changes in the presence of each DUB (Fig. 1A). Altogether, we profiled 30 DUBs in ten independent experiments. We defined candidate DUB substrates as proteins that were reduced in abundance in the immunoprecipitate in the presence of a DUB compared to no DUB addition (log_2_ fold change < −0.5, *p-value<*0.05). Because UbVS is an inhibitor of cysteine protease DUBs, we tested members belonging to the following classes: USP (21), OTU (5), MJD (3) and MINDY (1) (Fig. 1B). We observed that DUBs reduced the abundance of specific proteins from the immunoprecipitate, as expected if the enzymes remove ubiquitin from the isolated proteins (Fig. 1C). Furthermore, we noted that addition of DUBs paradoxically increased the level of a smaller set of proteins (Fig. 1C), in a DUB-specific fashion, suggesting that DUB addition may also indirectly stimulate ubiquitylation of certain proteins, perhaps by stimulating the activity of ubiquitin ligases or enhancing the rate of ubiquitin recycling.

**Figure 1.**
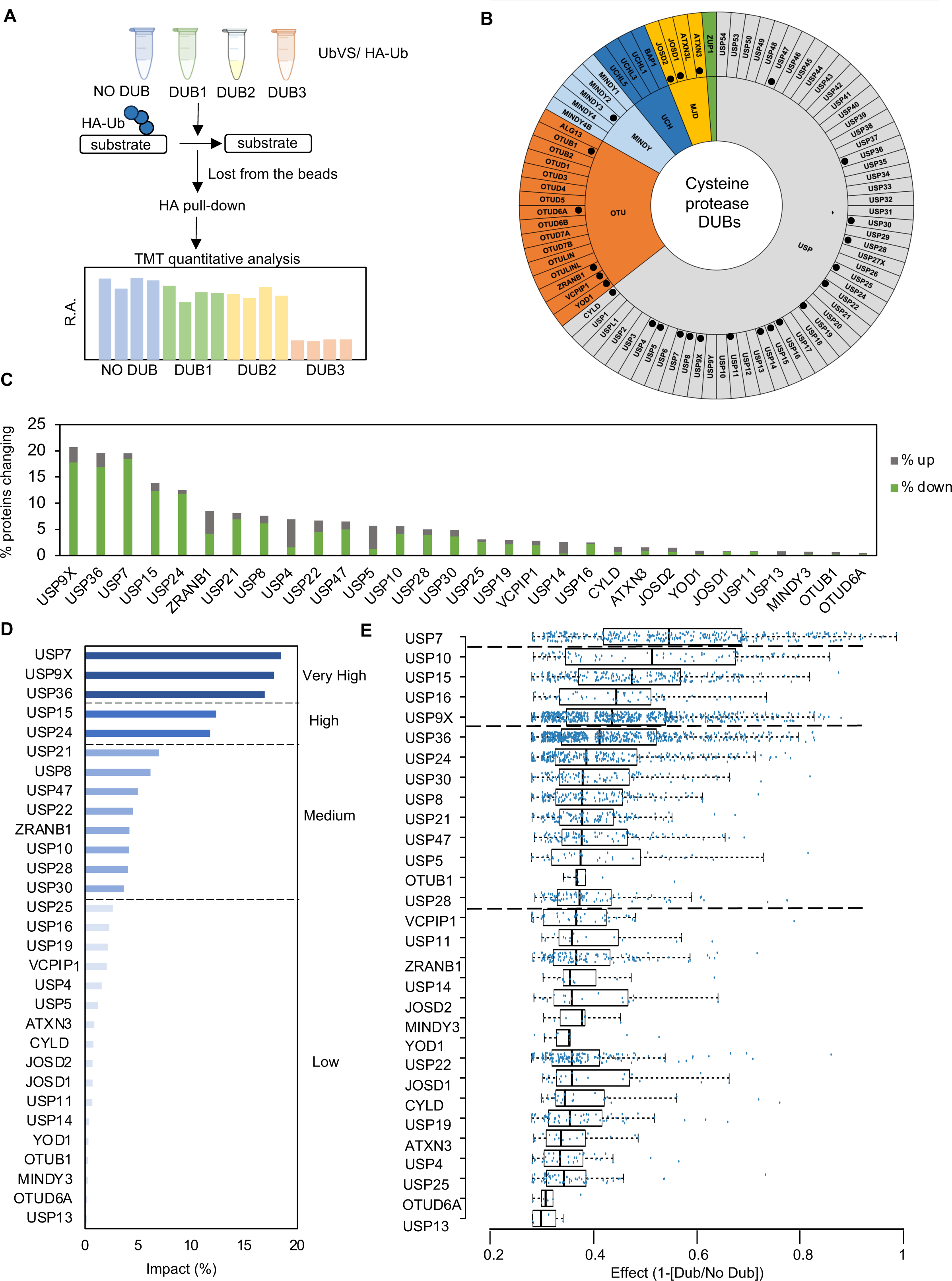
Summary of the approach and Impact and Effect of each DUB on the immunopurified proteins. **A)** *Xenopus* egg extract was incubated with 10 μM UbVS. After 30 minutes, individual recombinant human DUBs (800 nM) or no DUB (as control) were added in together with HA-ubiquitin (50 μM) and incubated for 30 minutes. HA-ubiquitin conjugates were immunopurified and characterized by an isobaric tag-based (TMT) mass spectrometry approach. Each experiment included four technical replicates for each DUB, with the exception of USP7, USP15 and USP21, which were tested with two technical replicates. **B)** Summary of the classes of cysteine protease DUBs. Black circles indicate the DUBs tested in our study. **C)** Percentage of proteins which changed in abundance the immunoprecipitate in the presence of the indicated DUB. **D)** Percentage of proteins decreasing from the beads (Impact) after addition of each DUB. **E)** Effect of the tested DUBs on their candidate substrates (Effect).

To compare the effects of the DUBs to each other, we calculated their Impact, defined as the percentage of proteins in the immunoprecipate that decreased after addition of a specific DUB. Impact varied substantially across the tested DUBs (Fig. 1D). Unsupervised clustering grouped the DUBs into four categories (Fig. 1D). We observed that Impact was higher for the USP DUBs compared to the non-USP DUBs. This finding is consistent with the fact that many USP DUBs have little ubiquitin-chain linkage specificity^16^, enabling them to act on a broader range of substrates and also remove all ubiquitin molecules attached to a substrate^20^. Impact showed a moderate correlation with the protein length of the DUBs (r=0.627) (Fig. S1B) as well as with the number of long disordered regions present in the DUBs (r=0.472) (Fig. S1C).

However, we also found that nine USP DUBs had low Impact (Fig. 1D). All of these low Impact DUBs reacted with UbVS *in vitro* (data not shown) with the exception of USP13, which is reported to be UbVS insensitive^23^, suggesting that they were catalytically active. We also compared how much each DUB reduced the levels of the immunoprecipitated proteins, which we refer to as Effect. The Effect presumably reflects the fraction of the substrate deubiquitylated by each DUB. Impact and Effect showed a moderate positive correlation (r=0.67) suggesting that DUBs with higher Impact tend to have a stronger Effect on their substrates (Fig. S1A).

We expected that high Impact DUBs might have more reported substrates in the literature. We therefore compared our results with a study that catalogued previously identified DUB substrates^10^. Both Impact and Effect showed a moderate correlation (r=0.484 and r=0.55, respectively) with the number of DUB substrates compiled by Elu et al.^10^ (Fig. S1D, S1E). The DUBs with the greatest number of reported substrates were USP7 (112), USP9X (65) and USP14 (79). Both USP7 and USP9X showed very high Impact in our screen whereas USP14 had a low Impact. Only a limited number of substrates have been reported for USP36 (6), USP15 (18) and USP24 (3), all of which had a high Impact in our study. Therefore, our approach has identified several DUBs with a broad activity that has not been recognized, perhaps because they have not been as well studied.

### Functional classification of the common set of immunopurified proteins

To characterize the functional roles of the immunopurified proteins and their relative sensitivity to DUBs, we identified a common set of 968 proteins that were reproducibly immunopurified in all experiments (Fig. S2A-B). We found that 374 of these proteins were candidate substrate of at least one DUB. We verified that restriction of the analysis to this set of proteins would generate a pool of candidate substrates representative of the entire data set. Indeed, the recalculated Impact of each DUB based on its activity on the common set of proteins was highly correlated with the original Impact (Fig. 1D) (r=0.984) (Fig. S2C). Next, we assigned the common set of proteins to 34 functional classes and calculated the percentage of proteins in each class that was DUB-sensitive (Fig. 2A). This percentage varied markedly. Two classes, centriolar/centrosomal proteins and heterogeneous nuclear proteins (HNRNPs) were highly sensitive to DUBs (Fig. 2A). On the other hand, 4 classes were almost completely insensitive to DUBs, including metabolic enzymes, proteases/peptidases, tRNA synthetases, and UB/UBL E1 enzymes. Particularly notable was the insensitivity of the metabolic enzymes, since over a hundred were reproducibly isolated across the ten experiments (Fig. 2A). The remaining classes showed varied sensitivity to DUBs. To understand the factors driving varied sensitivity within a class, we examined the behavior of subsets of proteins within each class (Fig. S3). Of the ribosomal proteins, only small ribosomal subunit proteins (RPS) were DUB sensitive; the large ribosomal subunit proteins (RPL) were not (Fig. S3A). In the case of the proteasome, the 19S complex and its associated shuttle proteins tended to be DUB sensitive, whereas the 20S complex was mostly insensitive (Fig. S3B). In evaluating E3s, we observed that RING and HECT ligases were sensitive, whereas CULLIN and most UBR ligases were insensitive (Fig. S3C). For proteins involved in folding, the DNAJ proteins and peptidyl-prolyl isomerases (PPIs) were sensitive, whereas members of the CCT complex and HSP class were insensitive. Altogether, this analysis indicates that the candidate substrates identified in the common set represent a wide variety of different protein classes with highly variable sensitivity to DUBs.

**Figure 2.**
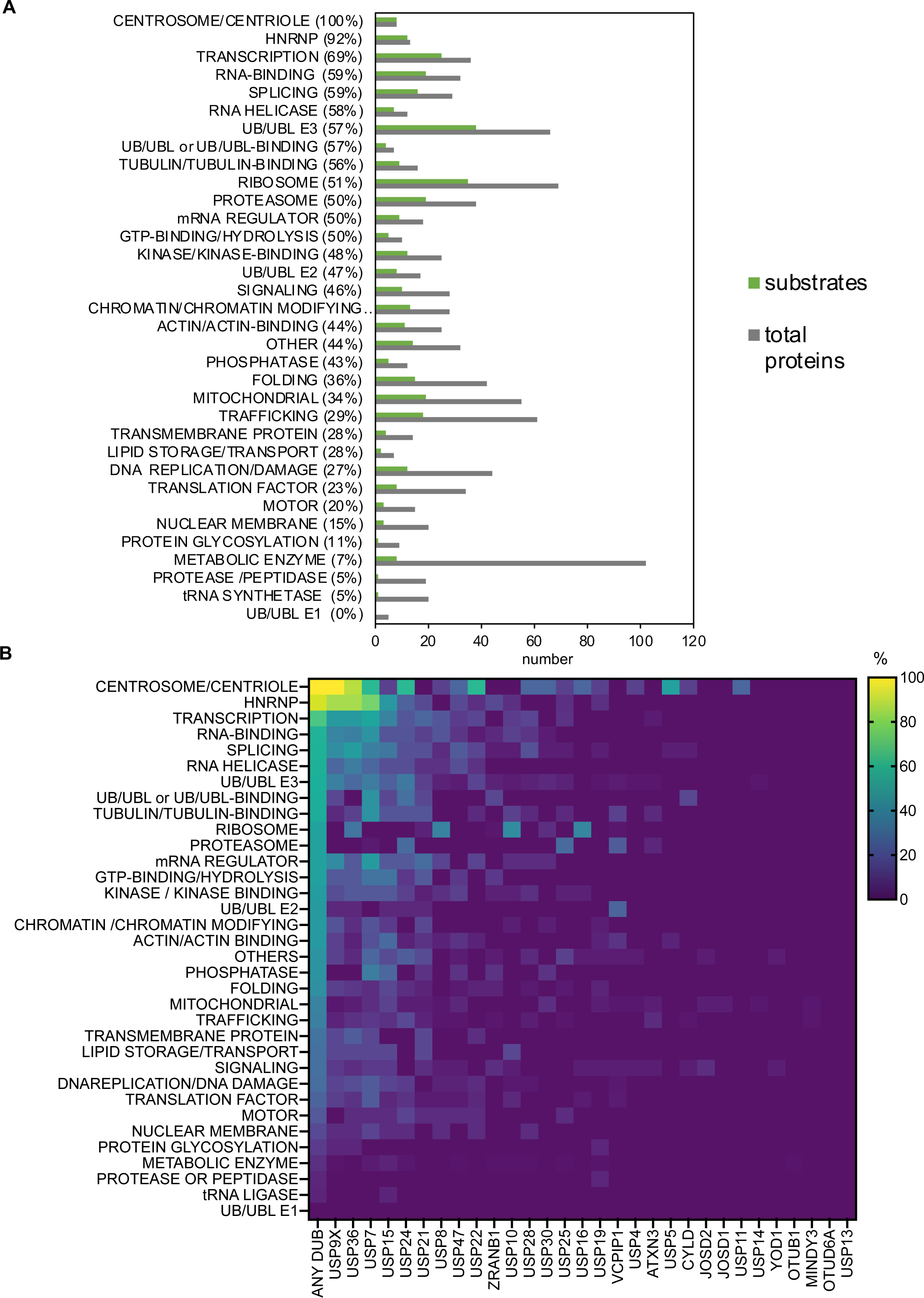
Functional classification of the common set of proteins and analysis of their DUB sensitivity. **A)** Number of proteins identified and number of candidate DUB substrates for each functional class. **B)** The heatmap reports the percentage of proteins in each functional class that are candidate substrates for any DUB (first column; arranged from highest to lowest) and for each DUB (all subsequent columns).

Next, we examined the sensitivity of each protein class to each DUB (Fig. 2B). We observed that the five DUBs with the highest Impact tended to share substrates belonging to the same functional classes. In particular, they acted on centrosome/centriolar proteins, HNRNPs, proteins involved in transcription and splicing, RNA helicases and E3s. However, the high Impact DUBs had little or no effect on many other classes of proteins, including the ribosome and proteasome. Instead, a distinct set of DUBs showed specific activity on these protein classes. The ribosomal proteins were specifically sensitive to USP16, USP10 and USP8, whereas the 19S proteasome components were sensitive to USP25 and VCPIP1. Together these findings suggest that the high Impact DUBs have a broad and overlapping specificity, suggesting they could be function redundantly in a cellular context. Yet despite their broad activity, the high Impact DUBs seem unable to access major classes of ubiquitylated substrates such as the ribosome and the proteasome, which are sensitive to a distinct set of DUBs.

### Comparison of DUB activity reveals patterns of specificity and redundancy

To characterize patterns of DUB activity and specificity, we first asked how frequently each protein was acted upon by DUBs. We found that 124 of 374 candidate DUB substrates (33%) were reduced in the immunoprecipitate in the presence of only a single DUB (Fig. 3A), showing a high level of specificity for a particular DUB. However, for the remaining 66% of candidate substrates, at least two or more DUBs showed an effect, suggesting that DUBs also function redundantly in this system. For some substrates, redundancy reached very high levels, as 93 proteins were candidate substrates of 5 or more DUBs (Fig. 3A).

**Figure 3.**
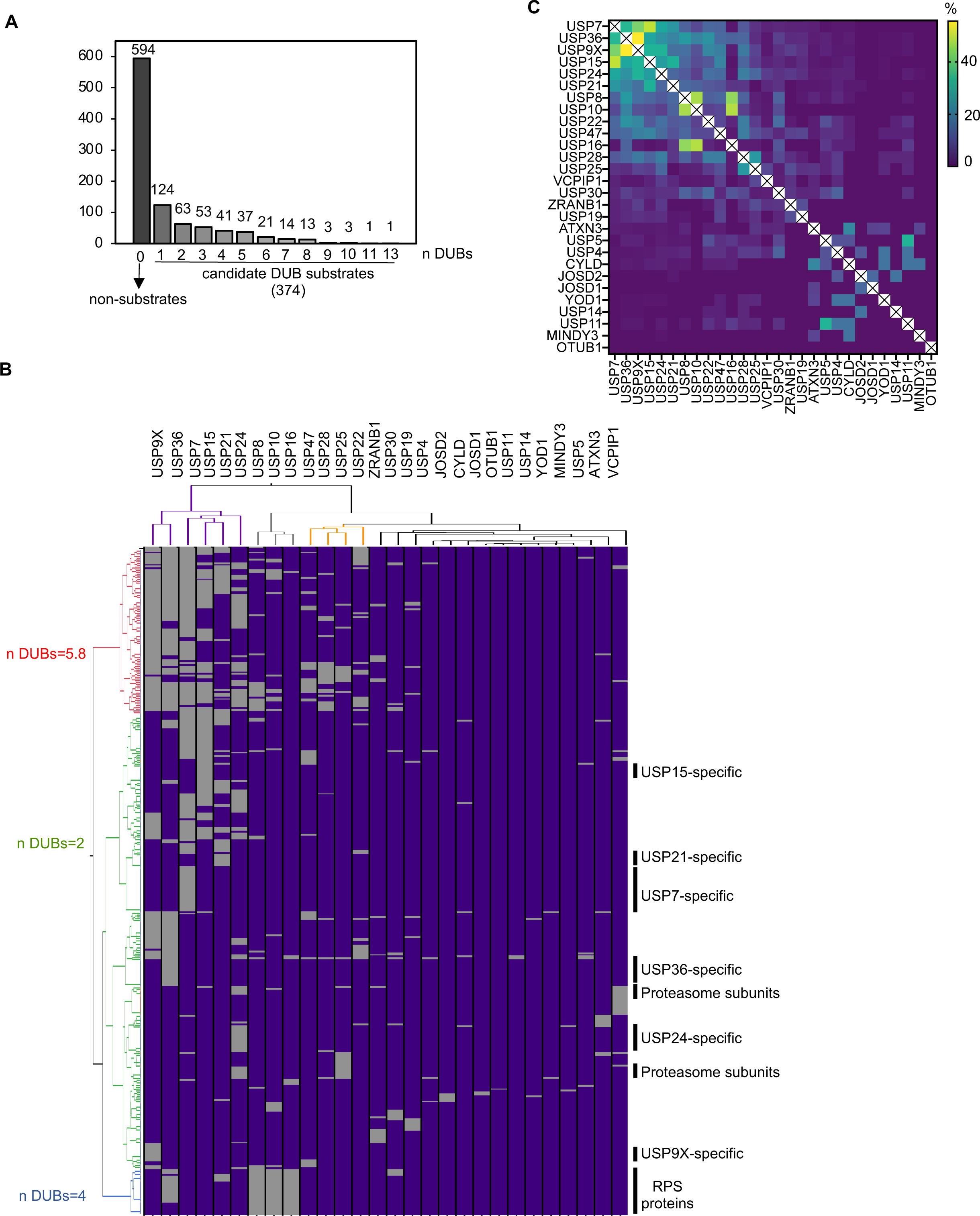
Analysis of the common set of ubiquitylated proteins reveals both specific and redundant patterns of DUB activity. **A)** The graph illustrates the number of proteins in the common set insensitive to DUBs (non-substrates) or sensitive to one or more DUBs (candidate substrates). **B)** Two-way hierarchical clustering analysis using Euclidean distance with Ward’s inter-cluster linkage based on the activity of each DUB against each candidate substrate. For the three main substrate clusters (rows; red, green and blue), the average number of DUBs per substrate is shown on the left. On the right, DUB-specific candidate substrates are labeled, as well as known functions of some of the proteins. **C)** The heatmap reports the percentage of shared candidate substrates between each pair of DUBs tested in our experiments.

We next performed an unsupervised two-way hierarchical clustering analysis based on the activity of each DUB against each candidate substrate. We found that the substrates fell into three main clusters (red, green, and blue) (Fig. 3B). Substrates in the red cluster were acted upon by many DUBs (mean=5.8). In contrast, substrates in the green cluster were acted upon by very few DUBs (mean=2) and included subclusters of proteins that were the targets of a single DUB. Finally, substrates in the blue cluster were composed entirely of the RPS proteins that were substrates of a small number of DUBs (mean=4).

In examining the clustering of the DUBs, we observed that the six highest Impact DUBs clustered together (Fig. 3B, purple DUB cluster) since they shared many substrates, particularly among the high-redundancy red cluster. However, these DUBs also possessed many unique substrates found in the green cluster (Fig. 3B). A second cluster of DUBs (gray) consisted of two medium Impact DUBs (USP8 and USP10) and one low Impact DUB (USP16), which shared a common set of substrates represented by the blue cluster of RPS proteins. Next, there were four other USP DUBs clustering together (orange DUB cluster) including three medium Impact DUBs (USP47, USP28, USP22) and one low Impact DUB (USP25). These four DUBs shared some substrates with the high Impact DUBs (red substrate cluster), but also possessed unique substrates not shared with the high Impact DUBs. The last cluster of DUBs (black) included all the low Impact DUBs and the two medium Impact DUBs ZRANB1 and USP30, whose substrates fell mostly into the green substrate cluster. The separate clustering of the high and low Impact DUBs reflects the fact that high Impact DUBs tend to act on different pools of proteins compared to the low Impact DUBs.

Our analysis showed that some DUBs act redundantly on a subset of their substrates. To quantify the degree of overlap, we determined the percent of common substrates for each DUB compared to others tested (Fig. 3C). USP9X and USP36 showed the highest fraction of shared candidate substrates, with USP7 and USP15 showing the second highest overlap. Furthermore, USP7 also shared many substrates with USP9X and USP36. In addition, there were other clear examples of high but specific redundancy, such as USP8, USP10 and USP16, which acted on the ribosomal proteins. USP28 and USP25 also shared a common set of substrates that were not shared by other DUBs. These enzymes are highly related to each other^24^, which may explain their common set of substrates. ATXN3 and CYLD, two low Impact DUBS, also showed a set of common substrates that were not affected by high Impact DUBs. These enzymes are known to act preferentially on K63-linked ubiquitin^22,25^, which may explain the overlap we observed.

### The presence of disorder in ubiquitylated proteins can favor substrate recognition, especially by high Impact DUBs

Our analysis revealed that high impact DUBs tend to act redundantly on a shared set of candidate substrates (Fig. 3B). However, these substrates belong to many different functional categories (Fig. 2B), raising the question of what features of these substrates render them sensitive to DUBs. Therefore, we asked if there were sequence characteristics or amino acid compositional bias that might explain the sensitivity of these proteins to DUBs. We found that that the average percentage of disorder for candidate substrates was higher than for the proteins insensitive to DUBs (Fig. 4A). We also found that substrates recognized by a greater number of DUBs showed an increased percentage of disorder (Fig. 4A). Furthermore, we observed that candidate substrates showed an increased frequency in the number of long-disordered regions (>50 amino acid) compared to non-substrates (Fig. 4B). Again, proteins that were candidate substrates of a greater number of DUBs showed the largest number of long disordered regions (Fig. 4B). These findings were also consistent with an analysis of amino acid composition (Fig. 4C), as candidate substrates were enriched, compared with non-substrates, in residues known to be present in disordered regions (especially R, G, S, P, and K) and depleted of hydrophobic amino acids that are present in non-disordered regions. Since the substrates of high Impact DUBs were more likely to recognized by multiple DUBs, we asked whether the substrates of high Impact DUBs were more likely to be disordered. Indeed, we found that the percentage of disorder of the substrates sensitive to at least one of the high Impact DUBs was higher compared to the substrates of non-high impact DUBs (Fig. S4A). Therefore, disorder may be important in explaining the ability of the high Impact DUBs to target a broad and overlapping set of substrates.

**Figure 4.**
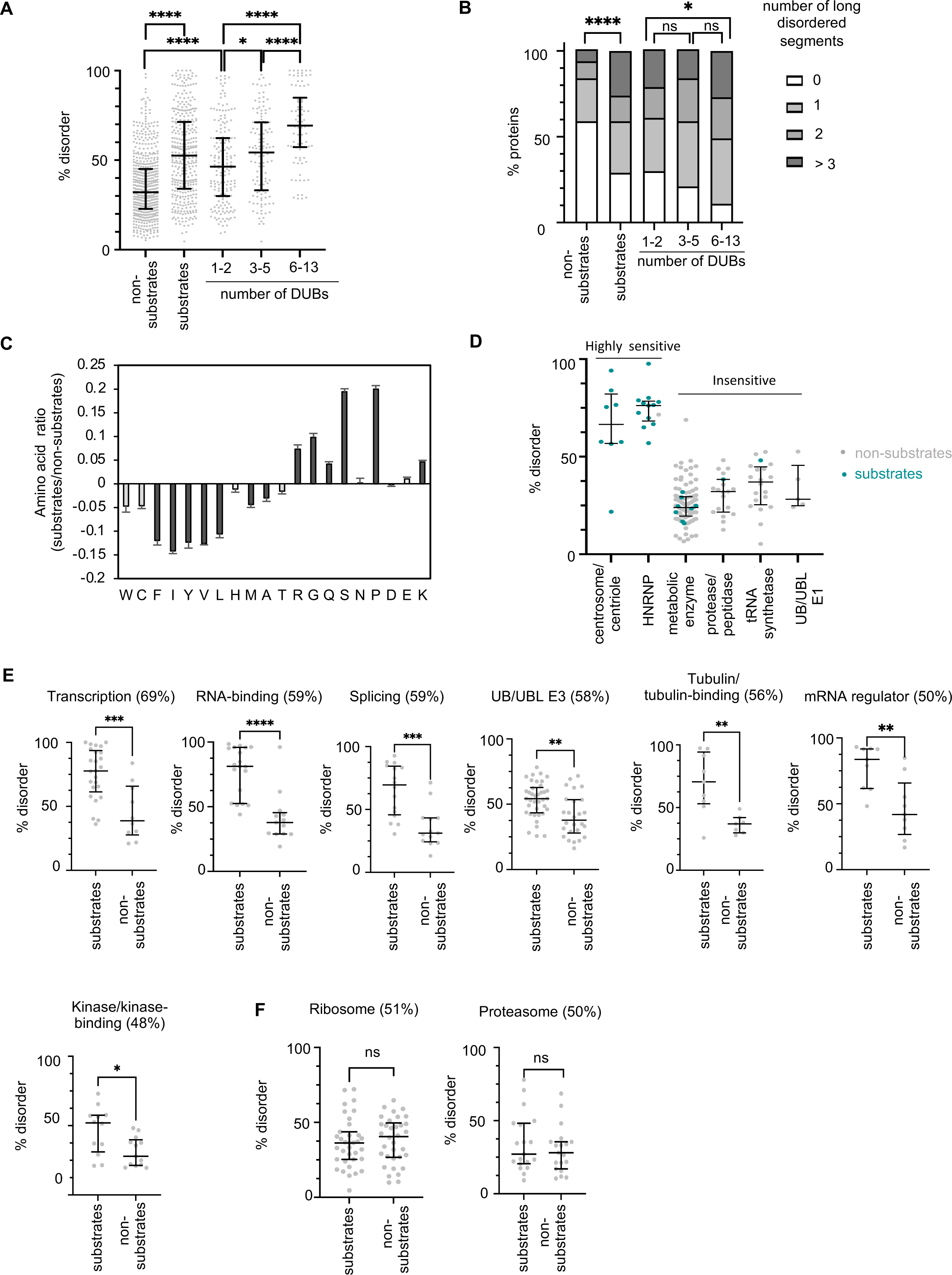
DUB substrates possess a higher percentage of disorder compared to non-substrates. **A)** Percentage of disorder and **(B)** number of long disordered segments possessed by the common set of proteins belonging to the indicated classes was calculated using the disorder prediction software Espritz^45^. Statistical significance was calculated using a two-tailed unpaired t test (A) or a Chi squared test (B). **C)** CompositionProfiler software^46^ was used to determine enrichment or depletion of amino acid frequency in substrates compared to non-substates. Error bar: standard deviation. Statistically significant changes (p-value<0.05, two sample t-test) are shown in dark gray. **D)** Percentage of disorder of proteins belonging to different functional classes highly sensitive or insensitive to DUBs. Substrates are colored in green while non-substrates are in grey. **E)** Functional classes of proteins in which candidate DUB substrates have a higher percentage of disorder compared to non-substrates. The percentage of proteins sensitive to DUBs is indicated on top of each graph. **F**) Percentage of disorder of proteasomal and ribosomal proteins. Statistical significance was calculated using a two-tailed unpaired t-test (**E** and **F**).

Next, we investigated whether the presence of disorder in specific functional categories of proteins correlated with their DUB sensitivity. We found that the two classes of proteins that were most sensitive to DUBs have a high percentage of disorder compared to the classes least sensitive to DUBs (Fig. 4D). Other classes showed a range of DUB sensitivities as well as a range of disorder. For many of these classes, candidate DUB substrates had a higher percentage of disorder compared to proteins insensitive to DUBs (Fig. 4E, Fig. S4B), especially in the categories that are the most sensitive to high Impact DUBs (Fig. 4E). However, several classes showed no difference in disorder between candidate DUB substrates and proteins insensitive to DUBs, suggesting that disorder was not a relevant feature for recognition of these substrates. This included ribosomal and proteasomal proteins that were insensitive to high Impact DUBs but sensitive to other classes of DUBs (Fig. 4F).

Our findings raised the possibility that previously reported substrates of high Impact DUBs might have an increased percentage of disorder. We analyzed the degree of disorder of known DUB substrates identified in human cells^10^ for USP7 and USP9X as they had the largest number of substrates^10^ and we compared the result to the low Impact DUB USP14 (Fig. S1E). We observed that USP9X and USP7 substrates had an increased level of disorder compared to the substrates identified for USP14 (Fig. S4C). Moreover, this higher degree of disorder was very similar to what we observed for the candidate substrates identified for USP7 and USP9X in our experiments (Fig. S4C). Therefore, an elevated degree of protein disorder seems to be a conserved feature of USP7 and USP9X substrates across human and *Xenopus* systems.

### Identification of high confidence candidate substrates of USP7, USP9X and USP36

To validate the reproducibility of our approach we performed a second independent experiment for the three DUBs that had the highest Impact in our screen (Fig. 1D): USP7, USP9X and USP36. We found that 41-58% of candidate substrates overlapped between the repeat experiments (Fig. 6A), with each DUB having a high Impact, comparable to what we observed in the initial experiments (Fig. 6B). The Effect for candidate substrates correlated moderately between the two experiments (Fig. 6C). We defined 389 proteins (116 for USP7, 275 for USP9X and 259 for USP36), that met both the *p-value* and fold-change threshold for both experiments, as high-confidence candidate substrates (Fig. 5A, S5A, S5B, S5C). We assigned these substrates to functional classes and calculated the percentage of substrates belonging to each class (Fig. 5D). We also confirmed that despite their broad activity, these DUBs do not act on many classes of proteins, including ribosomal and proteasomal proteins (Fig. 5D). Lastly, we compared the list of the high confidence substrates identified in our study with the substrates previously reported in the literature (Fig. S5A, S5B, S5C). We identified 46 previously reported substrates for USP7, 37 for USP9X, and one for USP36. This bias likely reflects the fact that USP7 and USP9X are much better studied than USP36. Overall, the degree overlap suggests that our method can reveal *bona fide* substrates that are conserved across experimental systems.

**Figure 5.**
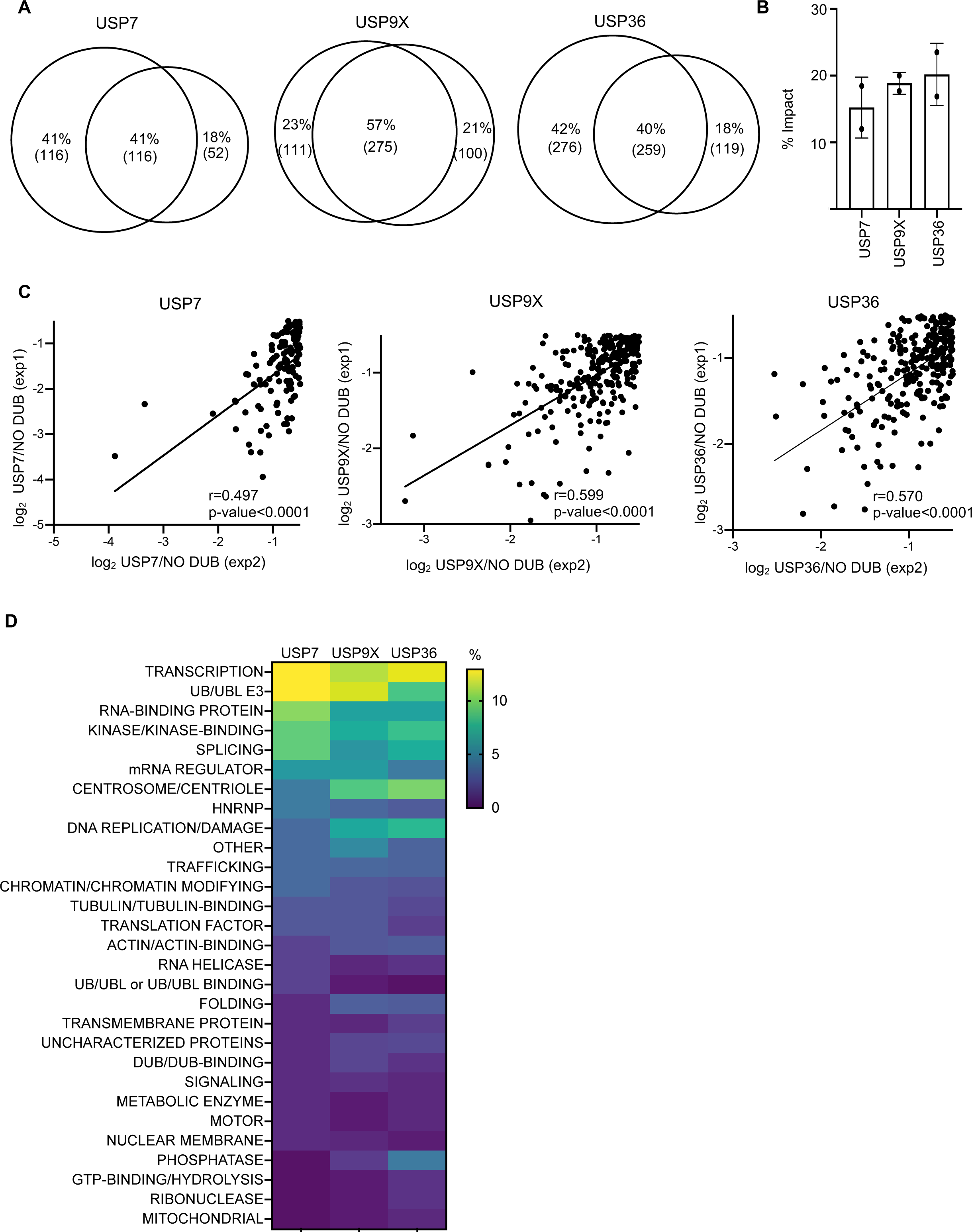
Analysis of the reproducibility of the approach for the three highest Impact DUBs: USP7, USP9X and USP36. **A)** The Venn diagrams illustrate the overlap of candidate substrates identified in two independent experiments for USP7, USP9X and USP36. **B)** Impact in the two independent experiments. **C)** Correlation between the Effect calculated in the two experiments (p-value < 0.0001; ****). Exp: experiment. **D)** The heatmap reports the percentage of proteins in each functional class that were identified as high confidence DUB substrates.

**Figure 6.**
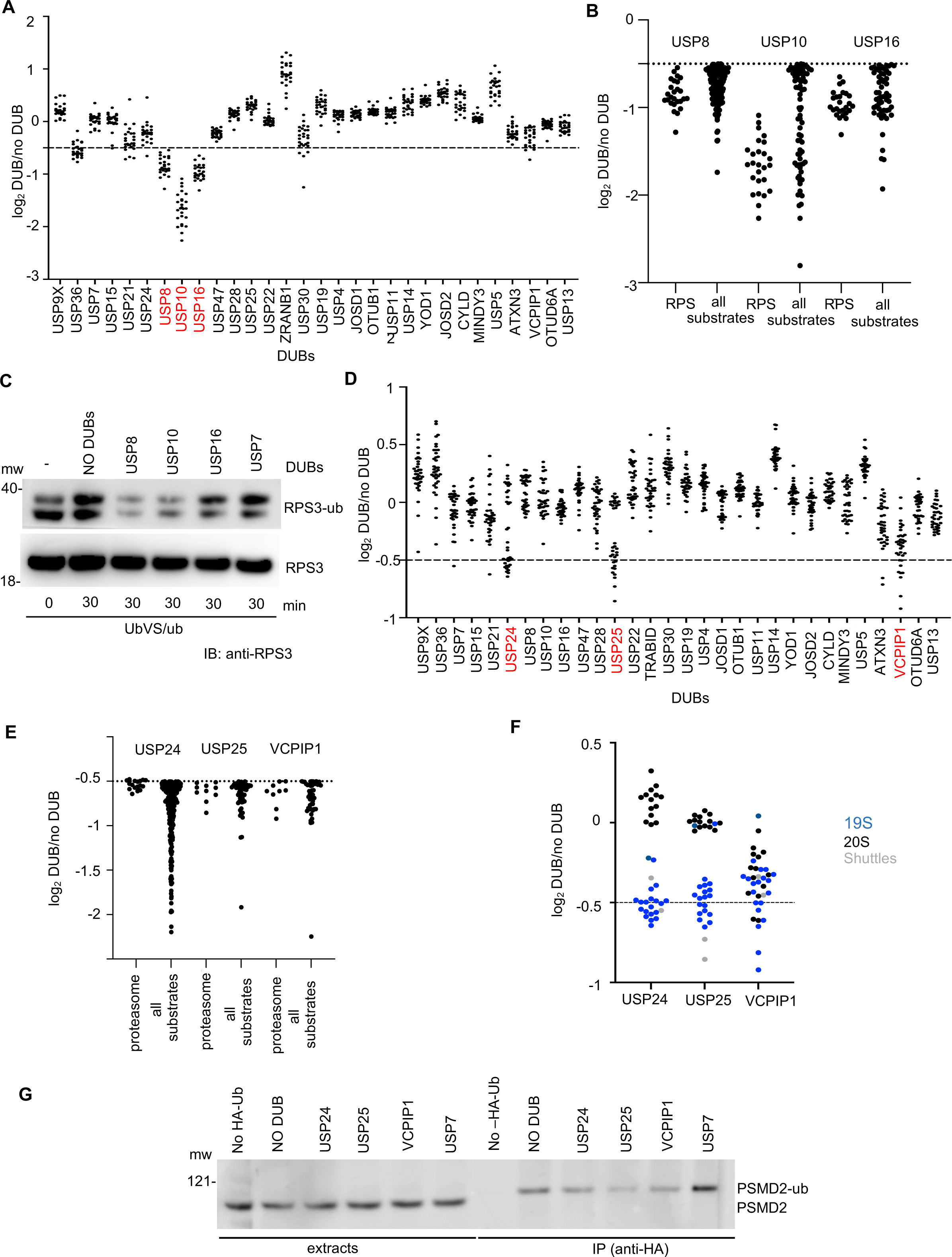
Two distinct sets of DUBs target ribosomal and proteasomal proteins. **A)** Profile of the ability of each DUB to alter the amount of immunoprecipitated RPS proteins, as measured in the original quantitative proteomic experiments, quantified by the log_2_-fold change in the presence or absence of each DUB (log_2_ DUB/No DUB). Each point represents a different RPS protein. Dashed line indicates the selection threshold for a candidate substrate (log_2_ DUB/NO DUB < −0.5). DUBs that caused a reduction in most RPS proteins are colored red. **B)** Comparison of the log_2_ fold change of USP10, USP8, USP16 on all their substrates compared to RPS subunits. **C)** Extract was treated with UbVS. After 30 minutes, ubiquitin and the indicated DUBs were added (time 0). Samples were collected at 30 minutes and processed for immunoblotting (IB) with antibodies against RPS3. **D)** Log_2_ fold change (DUB/NO DUB) for proteasome subunits. Dashed line: selected threshold (log2 < −0.5). DUBs that broadly caused decrease of proteasomal subunits from the immunopurified proteins are labeled in red. **E)** Comparison of the effect of USP24, USP25, VCPIP1 on their substrates and on the proteasomal proteins. **F)** Log_2_ DUB/NO DUB of USP25, USP24 and VCPIP on the different proteasome subcomplexes and on the proteasomal shuttle proteins. **G)** Effect of DUBs on ubiquitylation of the proteasomal subunit PSMD2. Extract was treated with 10 μM UbVS. After 30 minutes, HA-ubiquitin or buffer and the indicated DUBs (or buffer) were added (time 0). At 30 minutes, the HA-ubiquitylated proteins were immunopurified from each condition and samples were processed for immunoblotting.

### Ribosomal and proteasomal proteins are sensitive to two distinct sets of DUBs

Although the high Impact DUBs targeted a broad range of substrates belonging to many functional categories, they had little or no effect on many other classes of proteins. Some of these classes were insensitive to all the tested DUBs, such as the metabolic enzymes, whereas others were sensitive to a distinct set of DUBs, such as the ribosomal and the proteasomal proteins. Hierarchical clustering revealed that the ribosomal proteins (Fig. 3B, blue substrate cluster) were shared substrates of USP8, USP10 and USP16. (Fig. 2B). In particular, we found that 26 of the 30 candidate substrates shared between these three DUBs were RPS proteins (Fig. S6A). The RPL proteins were not affected (Fig. S6B). USP10 had the greatest Effect and showed a strong preference for RPS proteins relative to its other candidate substrates (Fig. 6B), as 9 out of 10 of the USP10 top candidate substrates were RPS proteins (Fig. S6C). USP16 showed a preference for RPS proteins as well, as 6 of its top 10 candidate substrates were RPS proteins (Fig. S6C). USP8 showed the weakest preference, with only one RPS protein among its top 10 candidate substrates.

Multiple studies have established that USP10 is a ribosomal deubiquitylase^26,27^, confirming the validity of our method. In addition, USP16 has been implicated in ribosomal biogenesis^28^, whereas USP8 has not been reported to deubiquitylate ribosomes. We therefore performed a second independent proteomic experiment for USP8 and USP16 and reproduced their effects on RPS proteins (Fig. S6D). We also observed a similar Impact in the repeat experiment (Fig. S6E). In comparing the substrates identified in the two independent experiments (Fig. S6F), we identified 62 and 20 high confidence substrates for USP8 and USP16, respectively (Fig. S6F, S7A, S7B). These high confidence substrates had limited overlap with the literature, likely because these two DUBs have a limited number of known substrates^10^. Lastly, we compared the high confidence substrates of the high Impact DUBs with those for USP8 and USP16 (Fig. S6G). Consistent with the results of the initial screen, we found limited overlap: USP8 shared 9% of its substrates with the high Impact DUBs and USP16 shared only 1%. Therefore, the high Impact DUBs and the ribosomal DUBs have largely non-overlapping substrate profiles in this system.

To further validate our findings, we analyzed the ubiquitylation status of RPS3 by immunoblot in extract treated with UbVS/ubiquitin after addition of USP10, USP16, or USP8, with USP7 as negative control. We chose RPS3 because it contained multiple UbVS-sensitive ubiquitylation sites in our previous study^13^ and is a known substrate of USP10^27^. We detected ubiquitylation of RPS3 that was UbVS-dependent and was reversed by addition of USP10 and USP8, with USP16 having only a minor effect (Fig. 6C), suggesting the latter enzyme may deubiquitylate another ribosomal protein. This is consistent with the fact that USP10 and USP16 regulate different aspects of ribosomal function^26,28^ and suggests that USP8 could be involved in the same pathway as USP10. Consistent with the results of our proteomic experiments, USP7 was not able to deubiquitylate RPS3.

A second protein complex that showed a unique profile of DUB sensitivity and was insensitive to high Impact DUBs was the proteasome. The proteasome has been shown to undergo reversible ubiquitylation^29^ and multiple DUBs are associated with it^30^, but no DUBs have been identified that affect its ubiquitylation. We found that USP25, VCPIP1 and USP24 were unique among DUBs in showing activity against the proteasome (Fig. 6D). USP25 and USP24 affected mostly components of the 19S proteasome and proteasome shuttle proteins, whereas VCPIP1 affected both 19S and 20S components (Fig. 6F). We validated this result (Fig. 6G) by treating the extract with UbVS/HA-Ub, adding USP24, VCPIP1, USP25 and USP7 as control and immunoprecipitating the HA-Ub-conjugated species. Immunoblot analysis of the 19S proteasome subunit PSMD2 showed that USP24, USP25 and VCPIP1 reduced the amount of ubiquitylated PSMD2 on the beads whereas USP7 did not, consistent with the results of our proteomic experiment.

### Confirmation of candidate DUB substrates with human orthologs

To further validate the findings of our proteomic experiments, we expressed human orthologs of candidate substrates in reticulocyte lysate and labeled them with ^35^S-methionine. Subsequently, we added the translated proteins to extract pre-treated with UbVS together with ubiquitin and the DUBs that scored in our screen. We selected three high confidence substrates of USP7, for which USP7 had a stronger effect relative to other DUBs: the splicing factor SMNDC1, the chromatin associated protein HMGB2 and the transcriptional coactivator of estrogen and progesterone receptors WBP2. UbVS treatment of the extract induced ubiquitylation of all three proteins confirming that they are acted upon DUBs in extract (Fig. 7A). Addition of USP7 caused almost complete deubiquitylation of these proteins (Fig. 7A, Fig. S8). USP21, which had little effect in the proteomic experiment, also had little effect in this experiment as well.

**Figure 7.**
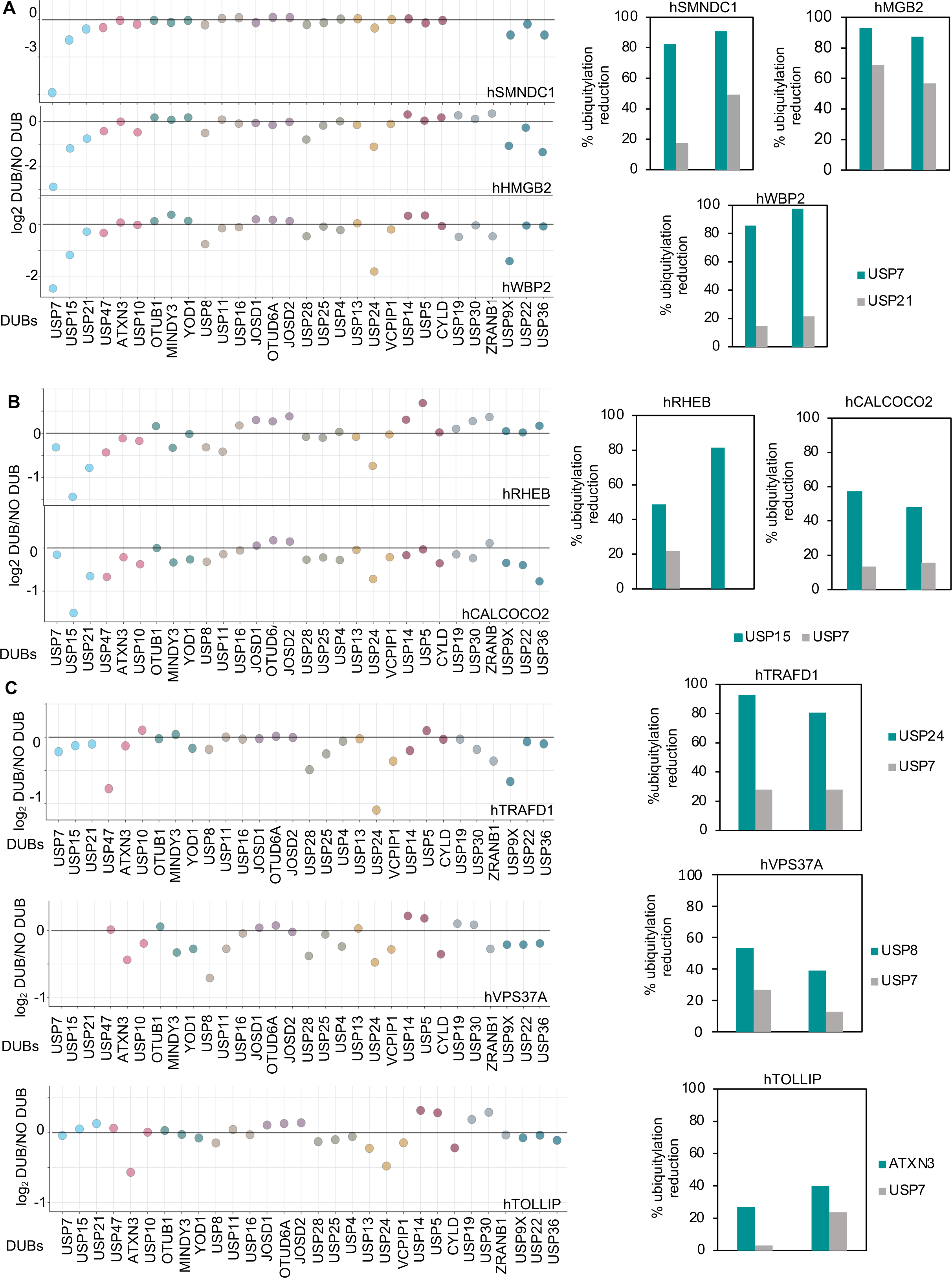
Independent validation of candidate DUB substrates. Left: Original quantitative proteomic data across the ten experiments for each candidate substrate selected for independent validation. Experiments are coded by color. Data points are missing if the protein was not identified in an experiment. Right: Effect of DUBs on recombinant human proteins. Candidate DUB substrates were translated in reticulocyte lysate and labeled with ^35^S-methionine. Labeled proteins were then added to extract pre-treated with UbVS together with HA-ubiquitin in the presence or absence of the indicated DUBs. After 30 minutes, samples were collected for SDS-PAGE and phosphorimaging. The percentage of reduction in ubiquitylation caused by addition of a specific DUB is reported for two independent experiments. The gels of the experiments are shown in Fig. S7.

Next, we chose candidate substrates that were sensitive to other high Impact DUBs, but insensitive to USP7. We tested the sensitivity of the autophagy receptor CALCOCO2 and of the mTORC1 activator RHEB to USP15 (Fig.7B) and the sensitivity of the TRAF-type zinc finger domain-containing protein1 (TRAFD1) to USP24. We observed that USP15 and USP24 decreased the ubiquitylation of CALCOCO2 and of TRAFD1 respectively, as we observed in our proteomic experiments (Fig. 7B and Fig.S8). Lastly, we tested candidate substrates of other DUBs such as the component of the ESCRT-I complex VPS37A, sensitive to USP8, and the autophagy protein TOLLIP, sensitive to ATXN3 (Fig. 7C, Fig. S8). For these substrates too, we detected an effect of USP8 and ATXN3 on the UbVS-dependent ubiquitylation of these proteins, consistent with the findings of the proteomic experiments.

## Discussion

By broadly inhibiting DUBs in *Xenopus* egg extract, we generated hundreds of endogenous ubiquitin-protein conjugates and developed a quantitative proteomic strategy to identify candidate substrates of 30 different DUBs. This method has the advantage of assessing DUB activity on many ubiquitylated proteins simultaneously, enabling a direct comparison of DUB activity and specificity. By adding back recombinant DUBs one-by-one in extract where broad DUB activity is inhibited, we also circumvented the challenges posed by DUB redundancy. Furthermore, since there is no ongoing transcription or translation in the extract, changes in levels of ubiquitylated proteins upon DUB addition are more likely to reflect direct effects of the added enzyme.

We observed that Impact was in general higher for the USP DUBs compared to the non-USP DUBs. This finding is consistent with the fact that USP DUBs have little ubiquitin linkage specificity, enabling them to act on a broader range of substrates^14,15,18,20^. Furthermore, USP DUBs may be more efficient at fully deubiquitylating their substrates^16,20^. The greater Impact of USP DUBs could also be due to the fact that they are active against both lysine and non-lysine ubiquitylation *in vitro*^17,18^. Finally, USP DUBs are, in general, larger enzymes than non-USP DUBs, which may provide more interaction domains for recognizing substrates.

However, we also observed some exceptions to the trend of USP DUBs showing higher impact than non-USP DUBs. For example, some USP DUBs showed lower impact. These included USP14, which acts mostly on substrates targeted to the proteasome and cannot remove the last ubiquitin from substrates,^31^ and USP5, which acts mainly on free ubiquitin chains^32^ rather than ubiquitylated protein substrates. The low impact of these DUBs in our assay is therefore consistent with their known mechanisms of action. Other examples include USP25 and CYLD, which are selective for K48^33^ and K63^34^ ubiquitin linkages, respectively. In contrast, some non-USP DUBs showed higher impact than other proteins in their class. For example, ZRANB1 and VCPIP1 are OTU DUBs with restricted ubiquitin chain linkage specificities^19^, yet they a had much greater Impact than other OTU DUBs. Both enzymes are larger than the other OTU DUBs tested, suggesting they may recognize their substrates using regions outside of their catalytic domain, which is usually sufficient for OTU DUBs to recognize their substrates^16^. Consistent with this, ZRANB1 has been shown to bind the ubiquitin ligase HECTD1 through one of its three Npl14 zinc finger domains^35^.

Among the most striking findings of our study are the substrate profiles of the high Impact DUBs (USP7, USP9X, USP36, USP15 and USP24), which share a broad and overlapping set of substrates that belong to many different functional categories. At the same time, the specificity of these DUBs is restricted, as they are unable to deubiquitylate large classes of proteins such as metabolic enzymes. Furthermore, they were not able to act upon substrates recognized by other DUBs, such as the ribosomal or the proteasomal proteins. By investigating if there were intrinsic substrate characteristics that could explain DUB specificity, we found that the presence of disorder was enriched among substrates in general, and among high impact DUB substrates in particular. Our analysis also revealed that substrates targeted by multiple DUBs have a higher percentage of disorder compared to substrates recognized by single and few DUBs, linking substrate disorder to DUB redundancy. How disorder facilitates recognition by these DUBs remains unclear. A previous paper suggests that USP36 may use its own unstructured regions to facilitate interactions with its substrates^36^. One possibility is that structural disorder facilitates sampling of weak protein-protein interactions that are critical for efficient substrate recognition, or the efficient positioning of the ubiquitin molecule into the DUB active site.

Our data show clearly that while disorder can favor recognition by high Impact DUBs, other DUBs can recognize distinct set substrates with a lower percentage of disorder. For example, USP8, USP10 and USP16 target the ribosome and USP25 and VCPIP1 act on the proteasome, protein complexes that show lower levels of disorder. Interestingly, these DUBs also tend to have a lower Impact, suggesting they use a distinct strategy for selecting their substrates compared to the high Impact DUBs. Furthermore, the known ubiquitin-chain selective DUBs showed a low impact in our study, consistent with the idea that they use distinct mechanisms for substrate selection. That we found many fewer candidate substrates for chain-selective DUBs suggests that ubiquitin is attached in the form of individual molecules or heterotypic chains rather than homotypic chains in *Xenopus* extract.

Although we identified fewer candidate substrates for the chain-specific DUBs, several of these proteins have roles consistent with the suggested biological function of these DUBs. For example, ATXN3, which preferentially targets K63-linked ubiquitin chains, is known to regulate autophagy and mitochondrial function^37,38^. Among the 16 candidate substrates we identified for this DUB, four were autophagy proteins (TOM1, TOLLIP, OPTN, EPS15) and four were mitochondrial proteins (PHB, PHB2, BIRC6, COX6B1). Indeed, we were able to recapitulate ATXN3-dependent deubiquitylation of TOLLIP with a radiolabeled human ortholog, validating our proteomic findings.

Even though our study revealed many candidate substrates, the majority (60%) of ubiquitylated proteins that we identified were in fact resistant to all the DUBs that we tested. Currently we do not understand the features that render these proteins resistant, other than that they are less likely to be disordered. It is possible that these proteins are substrates of DUBs we have not yet tested, or that they could be generally more resistant to the effects of DUB action. Another possibility is that multiple DUBs may need to act on these proteins to deubiquitylate them sufficiently to score in our assay.

Previously, we found that DUBs tend to act redundantly in controlling proteome stability in *Xenopus* extract^13^. Here we find that DUB redundancy is common for non-degradative substrates too. One reason that DUBs act redundantly could be that they deubiquitylate the same pool of substrates but in different cellular compartments. For example, in cells, USP7 localizes mainly to the nucleus^39^, USP9X to the cytoplasm^39^ and USP36 to the nucleolus ^40^. However, *Xenopus* egg extract lacks this compartmentalization, and therefore substrate recognition depends on intrinsic substrate characteristics rather than compartmentalization. We found that USP7, USP9X and USP36 are high Impact DUBs that recognize an overlapping set of substrates. In cells, this broad selectivity may be restricted by compartmentalization, in which each of these DUBs deubiquitylates a subset of this pool of substrates depending on their localization. Redundancy could also be a compensatory mechanism. This has been shown to be the case for USP24 and USP9X in the context of the survival of myeloma cells^41^. USP9X is highly expressed in myeloma cells and its inactivation reduces cell growth. In the absence of USP9X, USP24 is upregulated to sustain myeloma cell survival^41^.

Although several studies have employed proteomic approaches to study ubiquitylation in *Xenopus* extract^42,43^, this system has not been widely applied to study DUB function. Nevertheless, we observed many important overlaps with previous findings in mammalian cells, suggesting our findings are relevant to studies in other experimental systems. We observed that broad DUB inhibition, while having a limited effect on proteome stability^13^, causes accumulation of hundreds of ubiquitylated proteins in our system. These data suggest that the pool of non-degradative DUB substrates is larger than the pool of degradative substrates, which is consistent with a recent study in mammalian cells^44^. In fact, 48% of our common set of ubiquitylated proteins overlap with the proteins for which ubiquitylation increases after broad DUB inhibition in mammalian cells^44^, suggesting that the studies in the *Xenopus* system can reveal substrates conservated across different experimental systems. Furthermore, our identification of USP7 and USP9X as high Impact DUBs is consistent with the large number of substrates for these enzymes reported in the literature^10^. In fact, many of our high confidence candidate DUB substrates of these enzymes have been previously reported as substrates of the corresponding DUB in mammalian cells, including 46 of our USP7 substrates and 38 of our USP9X substrates.

Furthermore, both USP10 and USP16, which we found to act on the ribosome, have been previously implicated in ribosomal deubiquitylation^26–28^. In contrast, only a limited number of substrates have been reported in the literature for the other three high Impact DUBs we identified (USP36, USP24 and USP15), potentially because they are less well studied compared to USP7 and USP9X^10^. For USP36 the low number of substrates identified may be due to the essential nature of this DUB. Therefore, our approach has revealed several DUBs with broad activity that has not been recognized so far. Furthermore, our finding that USP7 and USP9X substrates are more likely to be disordered is also true for the reported substrates of these DUBs in human cells, suggesting this feature of these substrates is conserved across systems.

Our study has identified many candidate DUB substrates and provides insights into DUB specificity, but our study has also several limitations. Our approach may require that HA-Ub be fully removed from proteins to score, decreasing the sensitivity of our system to enzymes that partially deubiquitylate substrates. Moreover, because the HA-tagged ubiquitin cannot be incorporated in linear ubiquitin chains, proteins targeted by linear ubiquitin are missing from our analysis. It is possible that some DUBs may not be fully active when added as single enzymes to extract if they require specific posttranslational modifications or protein interactors that are missing in our system. It is also possible that recognition of some substrates by DUBs requires cellular compartmentalization that is absent in the extract system.

Altogether, our study presents a method that allows direct comparison of DUB activity against many endogenously ubiquitylated proteins. By examining the activity of single DUBs in a background of broad DUB inhibition, we can reveal the activity of DUBs on substrates that might otherwise be masked by functional redundancy. Our work highlights the importance of intrinsic protein substrate characteristics, such as the presence of disorder, for DUB recognition. Future work will be needed to determine how other protein substrate characteristics influence DUB-substrate recognition. We believe that for this purpose, the list of candidate DUB substrates that we generated in this study will be a valuable resource.

## Acknowledgments

We thank JW. Harper for the gift of plasmids (hORFeome collection) and the gift of the HA peptide. We thank D. Finley for the gift of USP21. We thank M. Wuhr for sharing the human mapping of the *Xenopus laevis* proteome 10.1. This work was supported by the Cell Biology Education and Fellowship Fund (VR) by the NIH grants R01 GM132129 (J.A.P.), R01 GM67945 (S.P.G), and R35 GM127032 (R.W.K.).

## Author contributions

V.R. designed, performed, and analyzed the experiments

V.R. prepared the samples for mass spectrometry with assistance from J.A.P.

J.A.P and S.P.G. performed TMT-mass spectrometric analysis

R.W.K. assisted with experimental design and interpretation

X.L. created the interactive web-based application

V.R. and R.W.K. conceived of the project and wrote the manuscript with input from other authors.

## Supplemental figure legends

**Figure S1.**
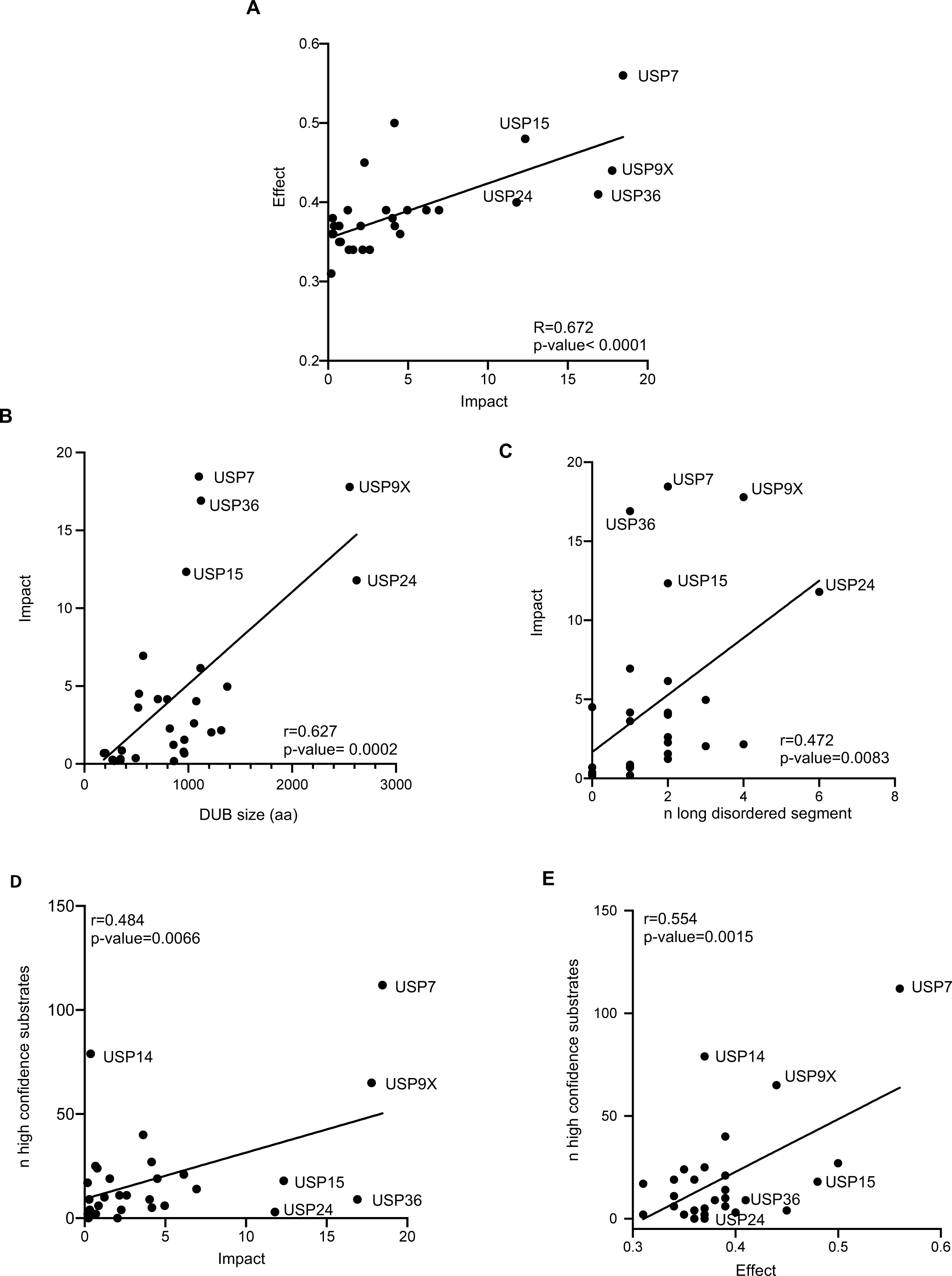
Correlation between Impact, Effect and intrinsic DUB characteristics. **A)** Correlation between Impact and Effect for the tested DUBs (p-value<0.0001; ****). **B)** Correlation between Impact and DUB size (p-value=0.0002; ***), aa: amino acid and number. **C)** Correlation between Impact and number of disordered segments longer than 50 amino acid (p-value=0.0083; **). **D), E)** Correlation between Impact (**D**; p-value=0.0066; **) or Effect (**E**; p-value=0.0015; **) and the number of high confidence candidate substrates identified in Elu et al.^10^.

**Figure S2.**
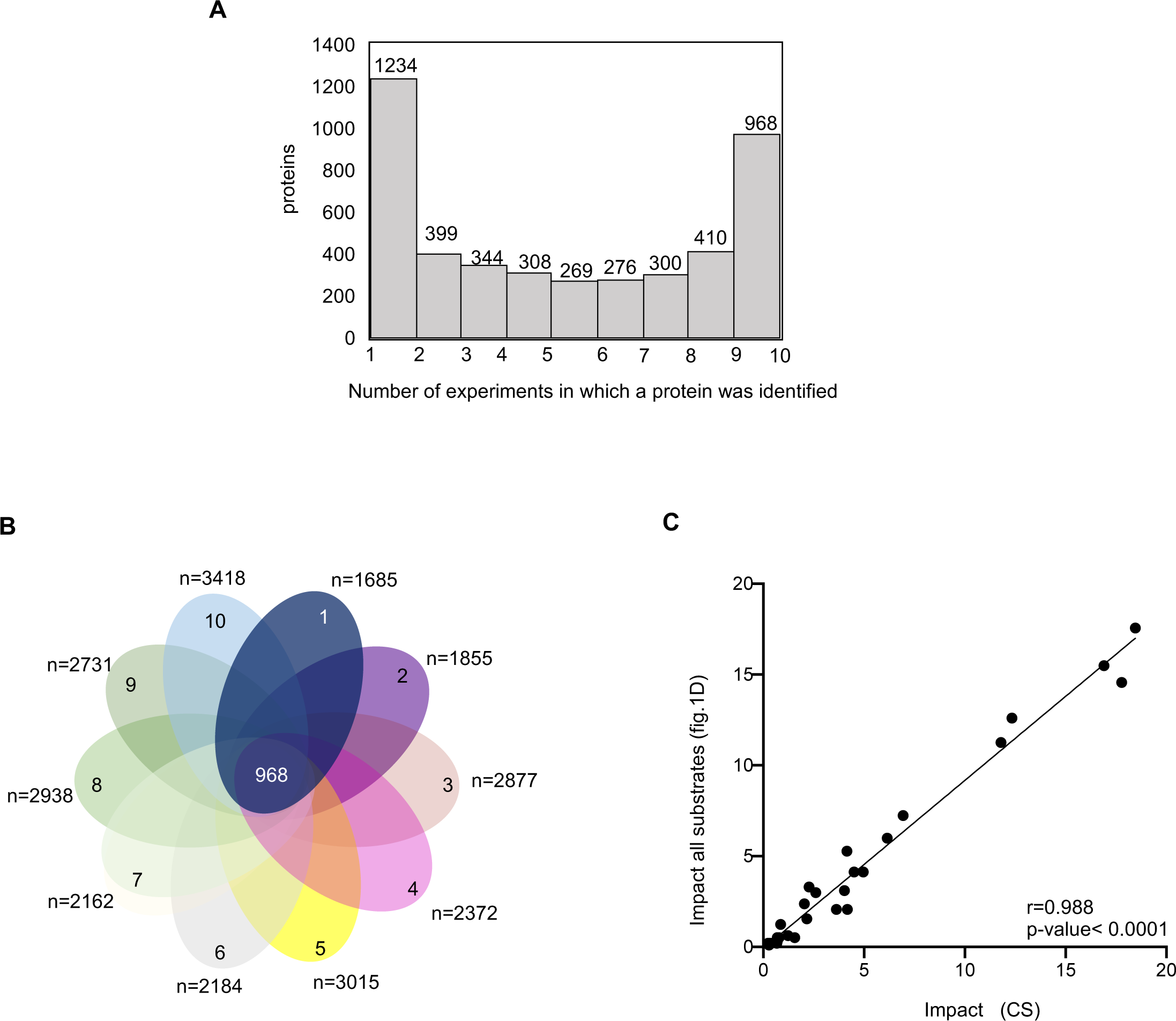
Summary of the frequency of protein identification across all experiments and definition of the common set of proteins (CS) **A)** The graph indicates the number of times in which a protein was identified across the 10 proteomic experiments. **B)** Definition of the common set (CS) of proteins as those 968 proteins immunopurified in all ten proteomic experiments. **C)** Correlation between the Impact of the DUBs calculated based on the candidate substrates identified in each single experiment (Fig. 1D) and the Impact calculated based on the candidate substrates of the CS of proteins (p-value<0.0001, ****).

**Figure. S3.**
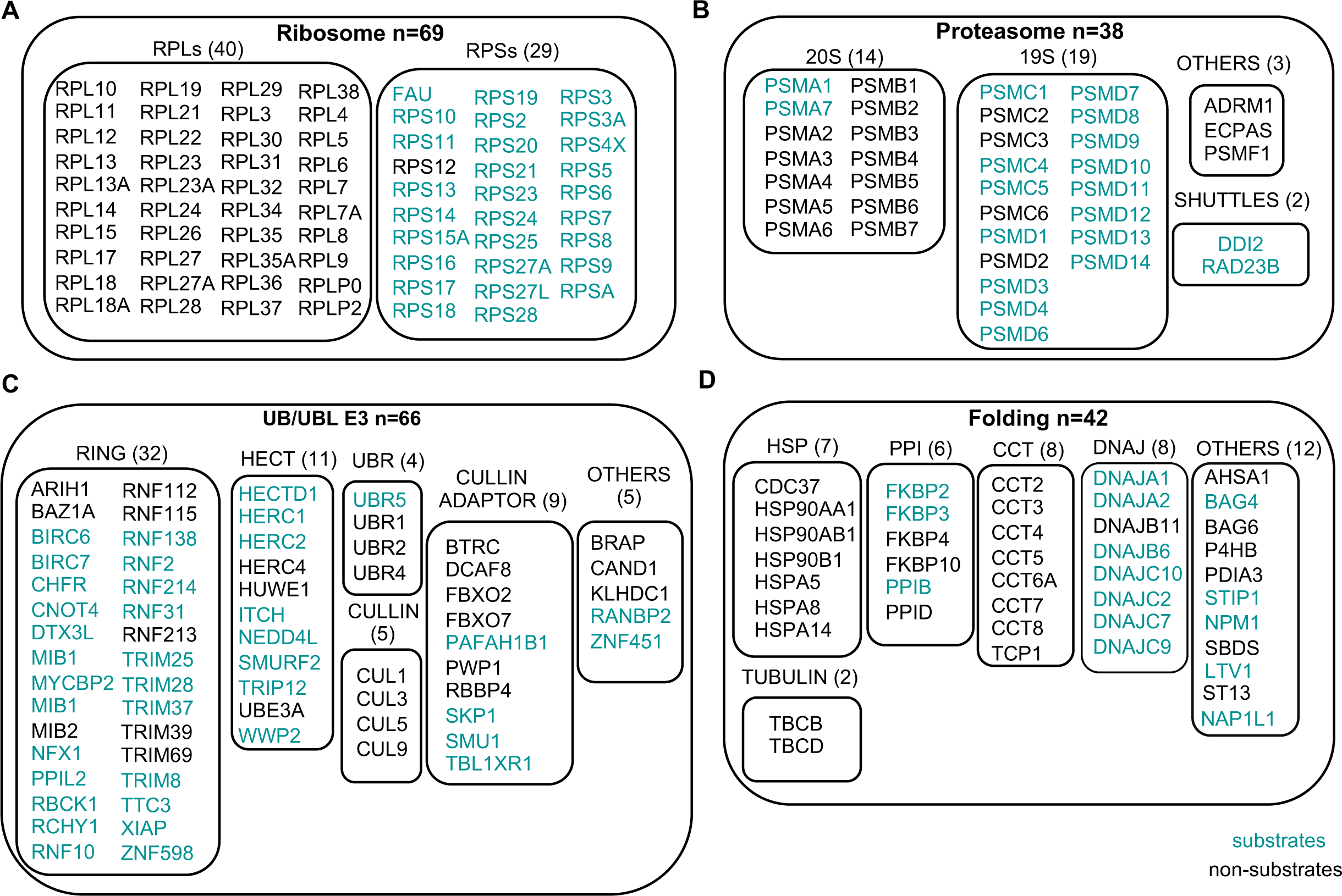
Sensitivity of different functional classes of proteins to DUB activity. **A)** Ribosomal proteins: large (RPL) and small (RPS) subunit proteins. **B)** Proteasomal proteins: 20S, 19S, shuttle proteins and other proteasome associated proteins. **C)** Ubiquitin ligase (UB/UBL E3) family proteins: RING, HECT, CULLIN, CULLIN adaptors and other ubiquitin/sumo ligases. **D)** Classes of folding factors: heat shock (HSP), prolyl isomerase (PPI), chaperonin-containing tailless complex polypeptide (CCT), DNAJ proteins, tubulin folding and other folding factors. Substrates are indicated in green, non-substrates in black.

**Figure S4.**
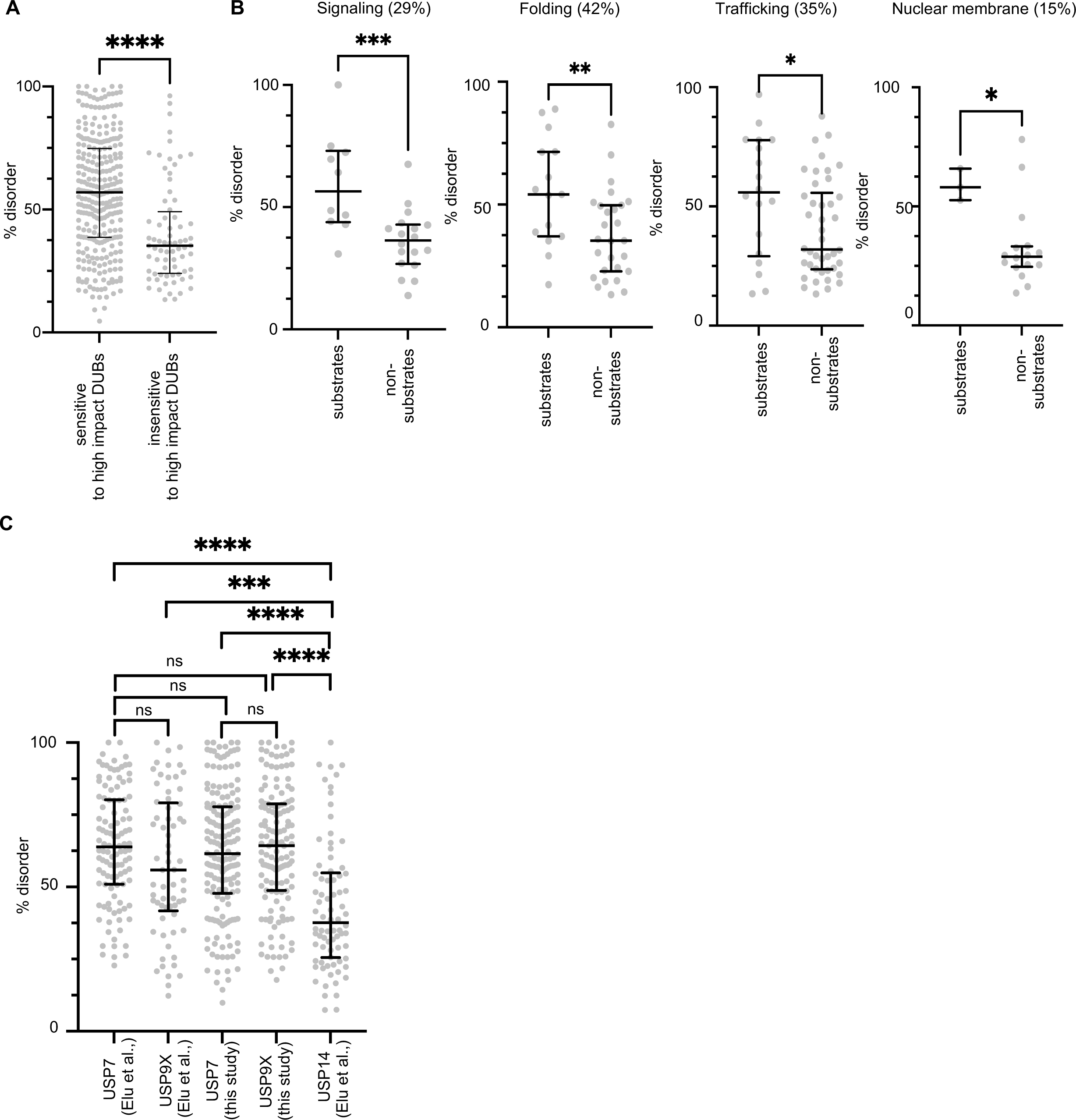
Classes of proteins in which disorder correlates with DUB sensitivity and disorder analysis of previously reported substrates in the literature. **A)** Percentage of disorder of candidate substrates of high Impact DUBs and substrates insensitive to high Impact DUBs but sensitive to other DUBs. Statistical significance was calculated using a two-tailed unpaired t test. **B)** Percentage of disorder of candidate substrates and non-substrates of the indicated classes of proteins. The percentage of proteins within the class that are DUB-sensitive is indicated at the top of the graph. Statistical significance was calculated using a two-tailed unpaired t test. **C)** Percentage of disorder of the substrates identified in our screen for USP7 and USP9X (common set of proteins) and of the substrates identified in the literature for these two DUBs and for the low Impact DUB USP14^10^. Statistical significance was calculated using a two-tailed unpaired t test.

**Figure S5.**
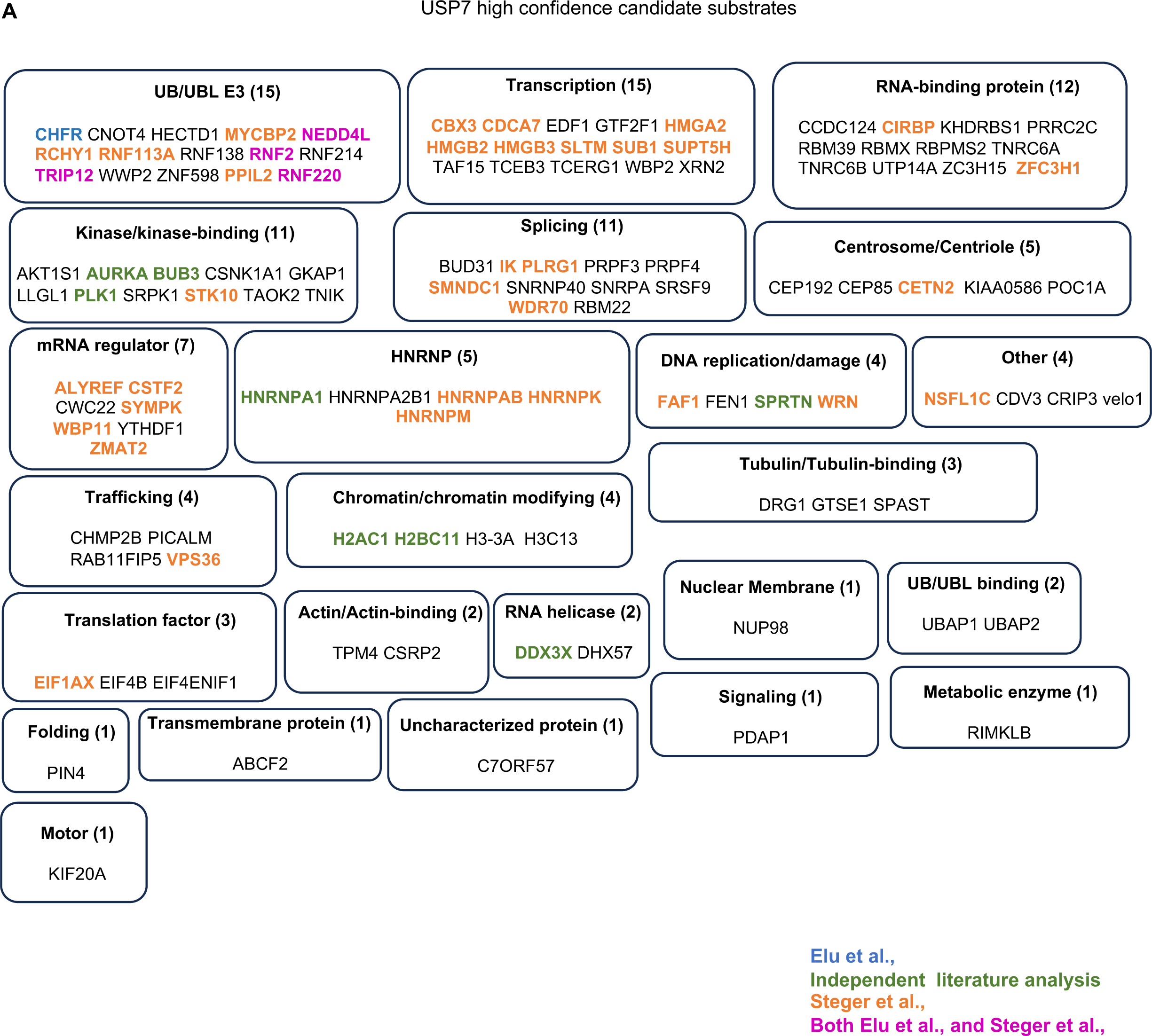
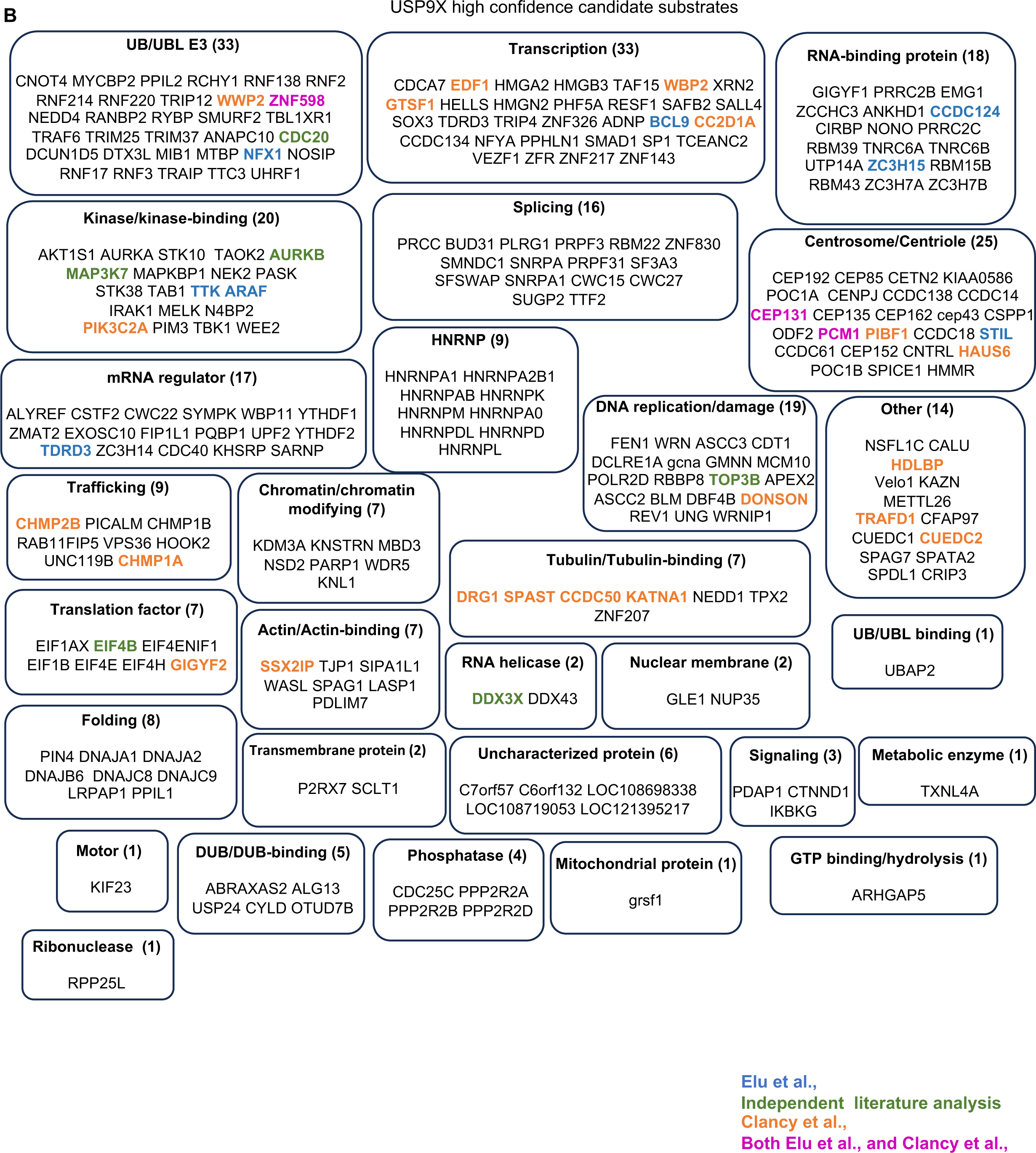
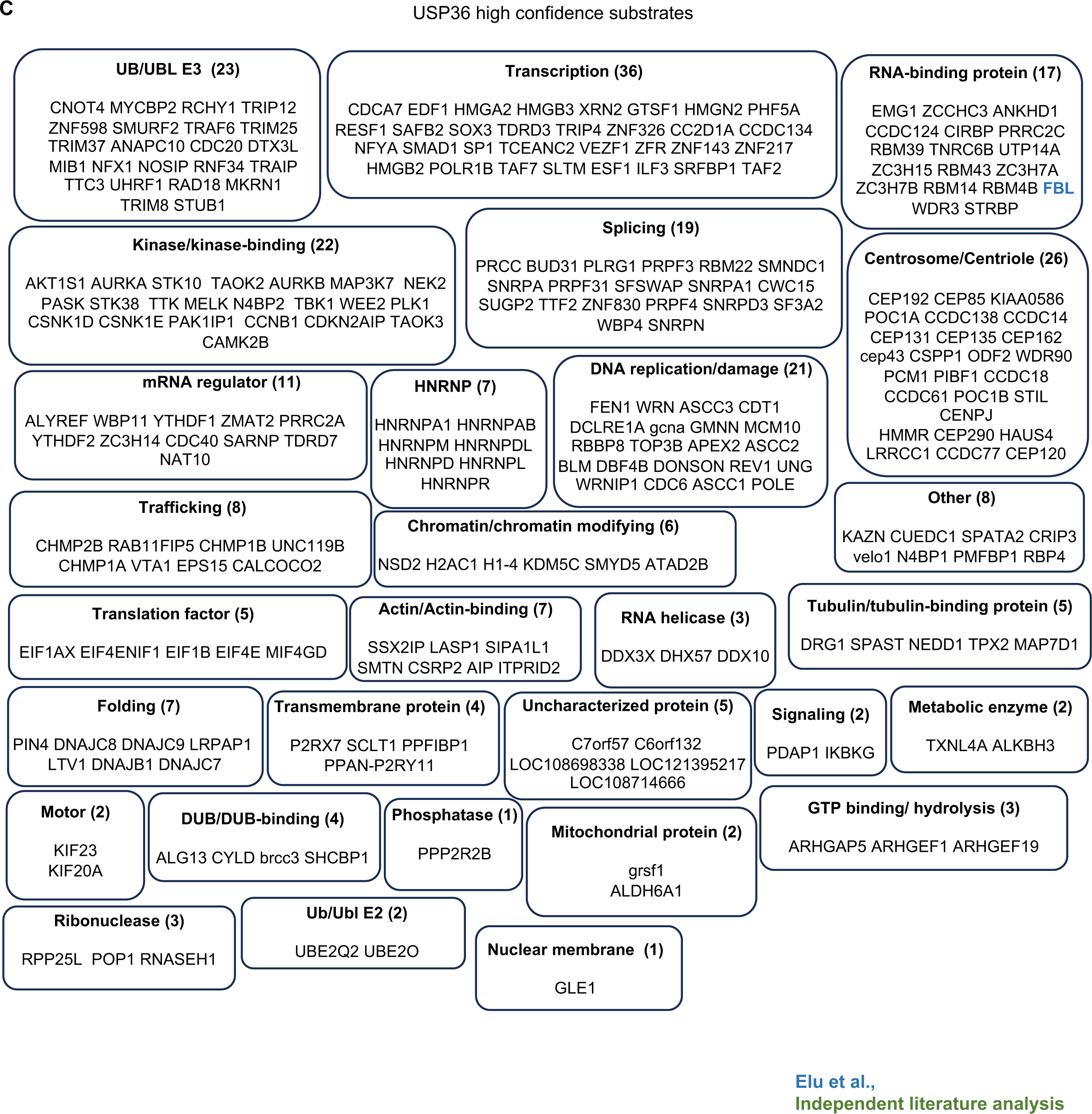
Functional classification of the high confidence candidate substrates identified for USP7, USP9X and USP36. List and functional classification of the high confidence candidate substrates identified for **(A)** USP7 **(B)** USP9X and **(C)** USP36. The number of candidate substrates belonging to each functional class is indicated in parenthesis. Substrates defined in the study of Elu et al.^10^ are in blue; substrates revealed by our independent analysis of the literature are in green. **A)** USP7 substrates identified in the work of Steger et al.^47^ (ubiquitylation sites significantly upregulated in presence of USP7 inhibitors compared to the control condition: log_2_ fold change > 1) are in orange and substrates identified by both Steger et al.^47^ and Elu et al.^10^ are shown in dark pink. **B)** USP9X substrates identified in Clancy et al.^48^ (log_2_ fold change < −0.05 change in protein stability in presence of the USP9X inhibitor compared to the control condition and p-value < 2 in presence of USP9X inhibitor) are in orange and substrates identified in both Elu et al.^10^ and Clancy et al.^48^ are in dark pink.

**Figure S6.**
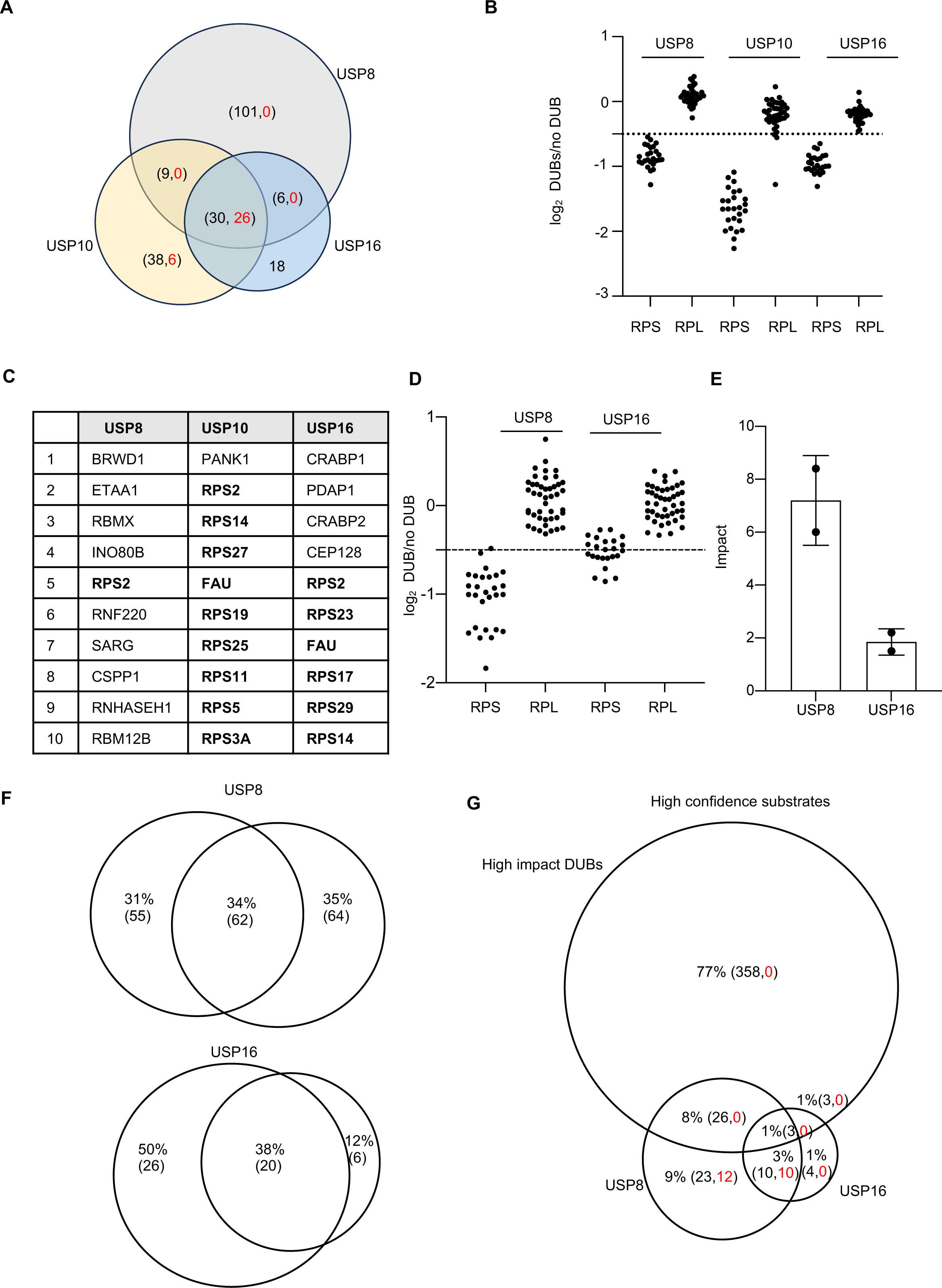
Effects of USP8, USP16 and USP10 on ribosomal proteins. **A)** Overlap of candidate substrates identified for USP8, USP10 and USP16 in the original screen. The number of RPS proteins is indicated in red. **B)** Comparison of the effects of USP8, USP16 and USP10 on the RPS (red) and RPL (black) ribosomal proteins. **C)** Top ten candidate substrates, based on log_2_-fold change, identified for USP8, USP16 and USP10. **D)** Effect of USP8 and USP16 on RPS and RPL proteins in a second independent experiment. **E)** Impact of USP8 and USP16 in two independent experiments. **F)** Overlap between substrates identified for USP16 and USP8 in two independent experiments. **G)** Overlap between the high confidence candidate substrates identified for the high Impact DUBs, USP8 and USP16. The number of RPS subunits is indicated in red.

**Figure S7.**
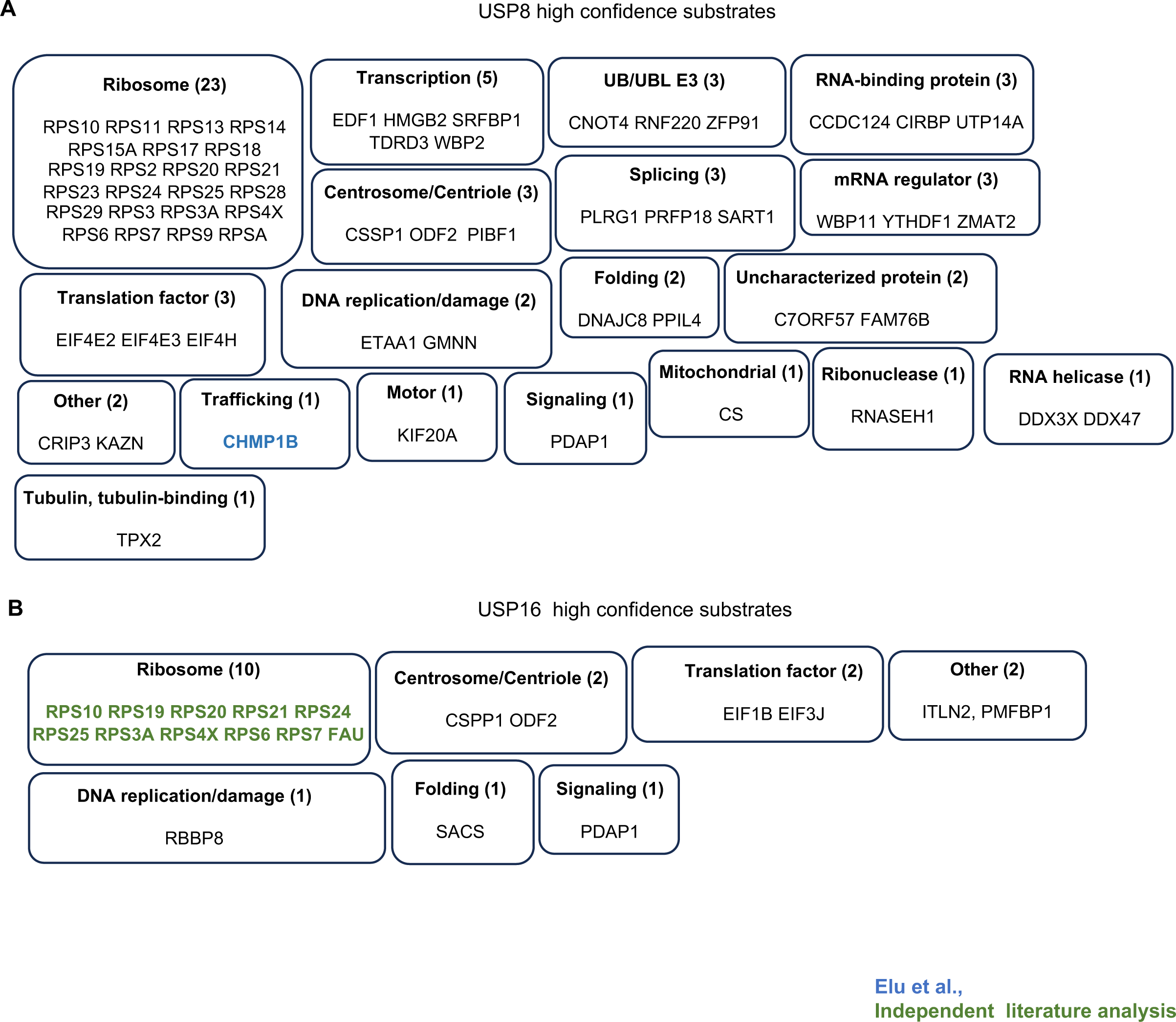
High confidence candidate substrates identified for USP8 and USP16. Functional classes of the high candidate substrates identified for USP8 **(A)** and USP16 **(B)**. The number of candidate substrates belonging to each functional class in indicated in parenthesis. Substrates identified in Elu et al.^10^ are in blue while substrates identified by our independent analysis of the literature are in green.

**Figure S8.**
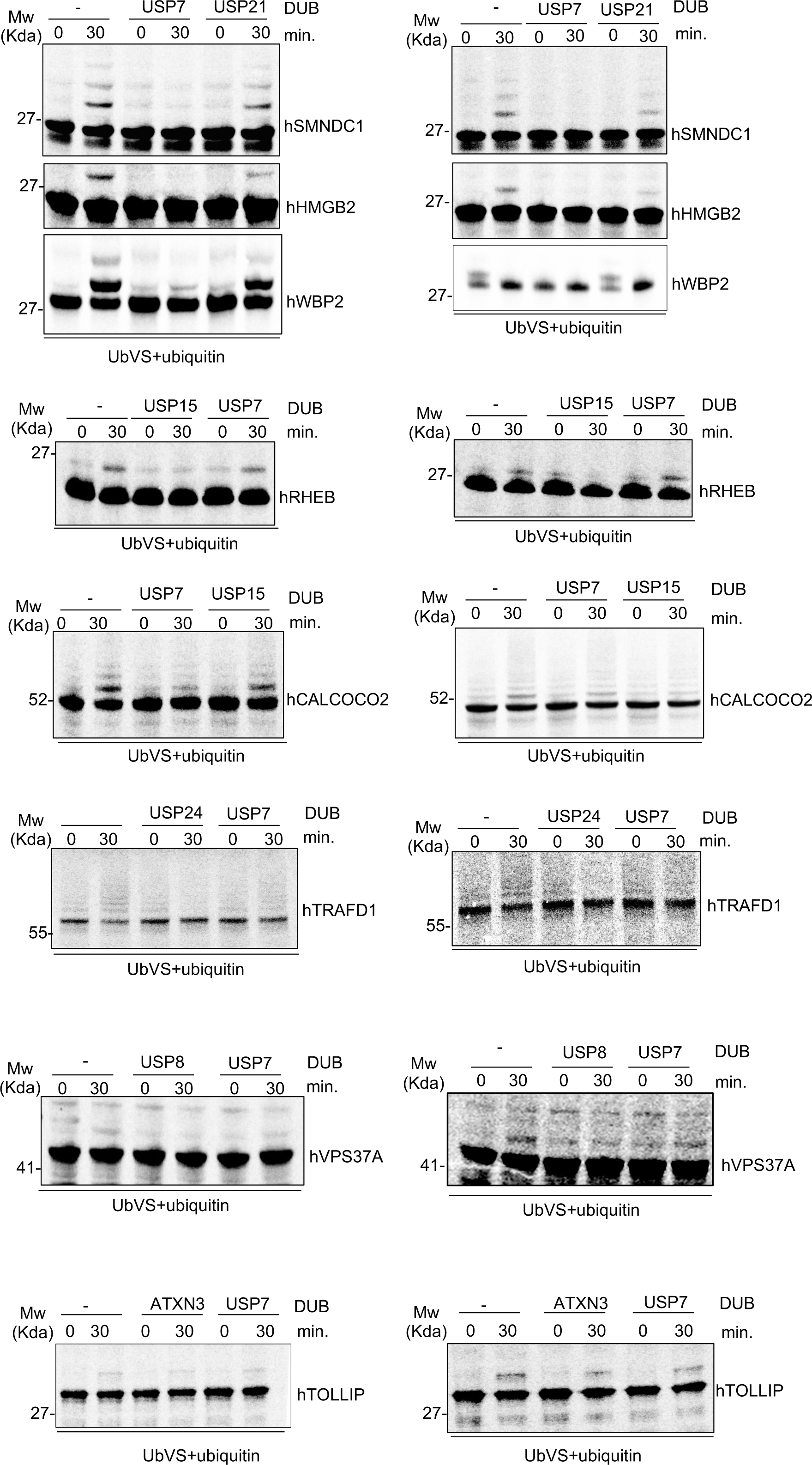
Images of the experiments quantified in Figure 7 are shown.

**Figure S9.**
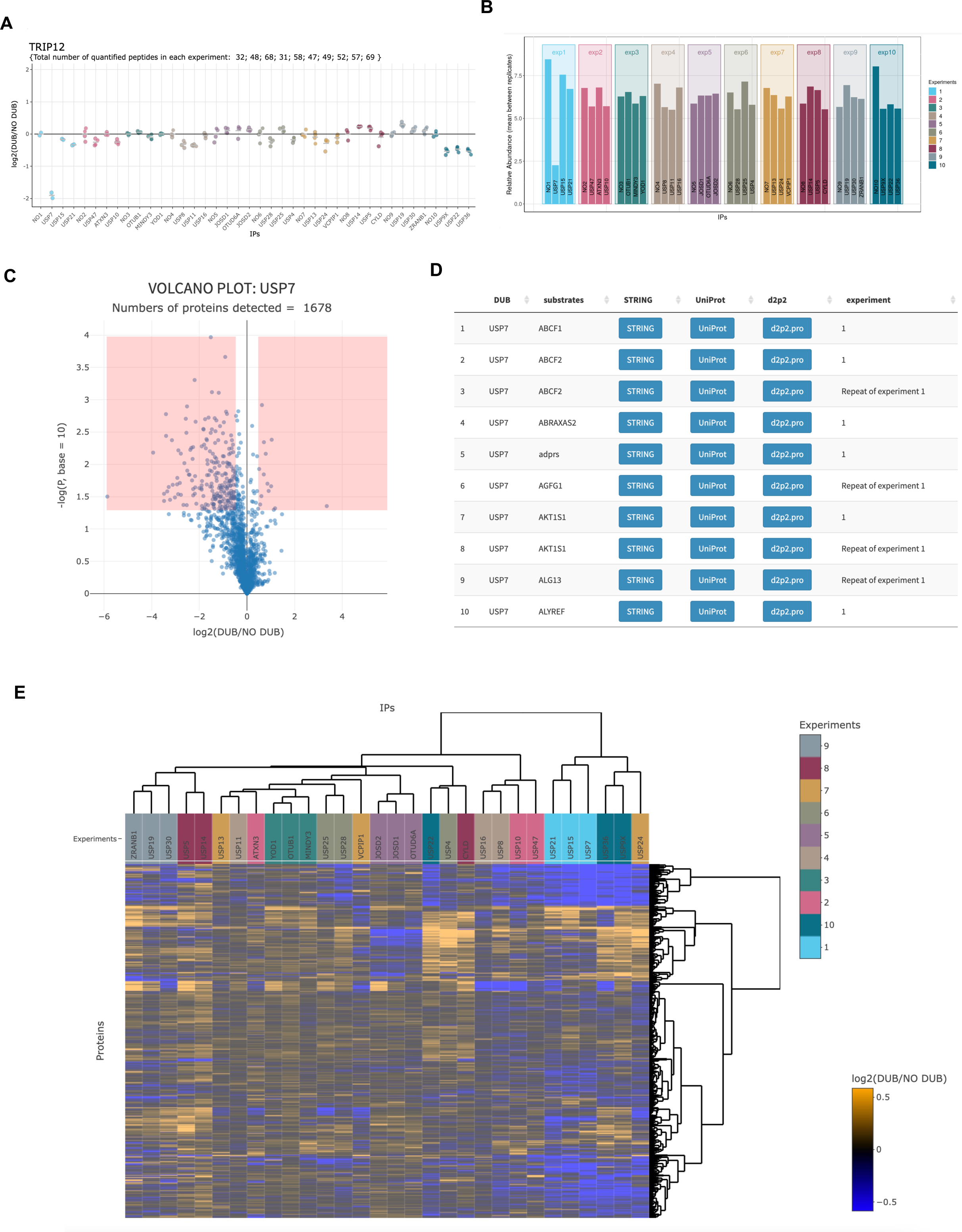
Description of the web-based application created to share our data. **A)** The profile reports the log_2_ fold change in protein abundance of TRIP12 in the immunopurified proteins after addition of each DUB compared to the NO DUB condition of the same experiment. The relative abundance profile (average between replicates) of TRIP12 is shown in **(B)**. TRIP12 is a known USP7 substrate^10^. **C)** Volcano Plot of TMT analysis comparing the proteins detected in the anti-HA immunoprecipitated in the presence or absence of USP7. Statistical significance (− log_10_ *p*-value) is plotted against ratio (average log_2_) of the samples. The red rectangles highlight the proteins that meet the cutoff for both log_2_ fold change and statistical significance. **D)** The table reports candidate substrates identified for USP7 together with links to websites that present additional useful information about the proteins. **E)** Hierarchical clustering of the log_2_ fold-change data for the common set of proteins is shown.

**Figure S10.**
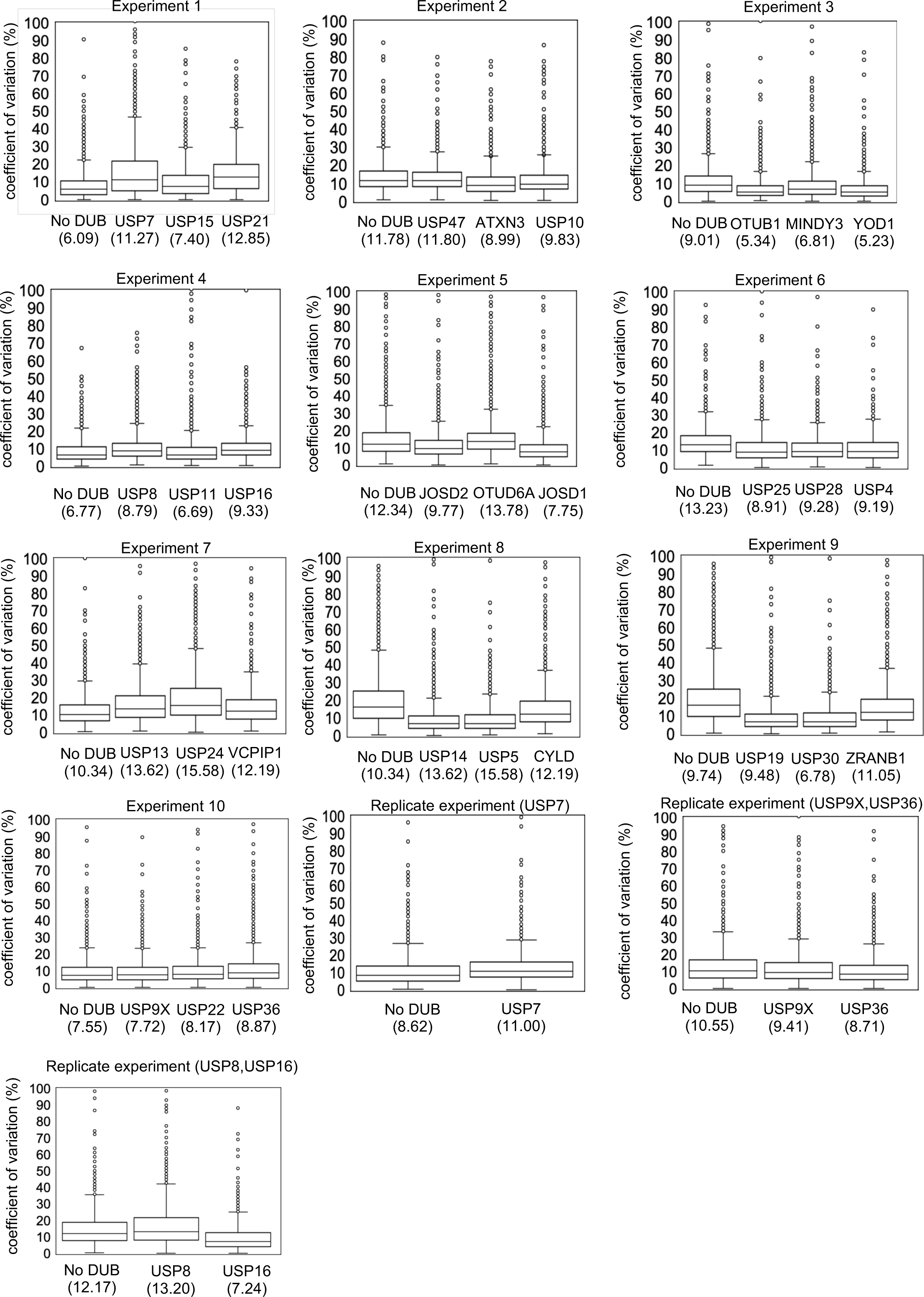
Coefficient of variation. Box plots showing the distribution of the coefficient of variation (CV), which is calculated by dividing the standard deviation of the peptide profiles by the mean, for the replicates at protein levels for all the proteomic experiments. Median values are reported under the graphs.

## STAR Methods

### Lead Contact

Further information and requests for reagents should be directed to the lead contact: Randall W. King

### Materials Availability

This study did not generate new unique reagents.

### Data and Code Availability

The mass spectrometry-based proteomics data have been deposited to the ProteomeXchange Consortium ^49^ via the PRIDE partner repository.

## EXPERIMENTAL MODEL

### Preparation of *Xenopus laevis* egg extract

Female *Xenopus laevis* (adult 3-5 years old) were used to produce the cytoplasmic egg extract. They were housed and maintained at Harvard Medical School. The extract was prepared as previously described^50^. All experiments involving animals were approved by the Harvard Medical School Institutional Animal Care and use Committee (IACUC) and conform to relevant regulatory standards (Protocol number IS117-6).

## METHOD DETAILS

### Gene nomenclature

The human gene or protein symbols have been used in the manuscript for simplicity and consistency. Multiple isoforms of the same *Xenopus* protein were grouped together under the corresponding human gene. Assignment of the human gene was based on the human mapping of the *Xenopus laevis* proteome 10.1 shared by M. Wuhr. This unpublished human mapping resource is the most updated. For the few cases in which the human gene was not assigned, the *Xenopus* gene names were used.

### Antibodies

The following antibodies were used for immunoblotting: anti-RPS3 (Abcam, ab140688), anti-PSMD2 (Fortis Lifescience, A303-854A), anti-rabbit IgG-HRP (NA934V) and anti-mouse IgG-HRP (NA931V) (GE Healthcare, GENA934 and GENA931).

### Experiments comparing DUB activity on HA-immunopurified proteins

*Xenopus* egg extract was treated with 10 μM UbVS (R&D SYSTEMS, U-202) for 30 minutes at 24°C. Extract was then divided in single reactions consisting of 20 μl each. Single human recombinant DUBs (800 nM) were added together with 50 μM human recombinant HA-tagged ubiquitin (R&D SYSTEMS, U110) to each reaction. The reactions were then incubated 30 minutes at 24°C. After that, the reaction was diluted three times with XB/high salt buffer (10 mM potassium HEPES pH 7.7, 500 mM KCl, 0.1 mM CaCl2, 1 mM MgCl_2_, 0.5% NP40 and 5mM NEM) and incubated with 15 μl of anti-HA agarose beads (Sigma Aldrich, A2095) rotating at 4°C (90 minutes). Beads were washed four times with 500 μl of XB/high salt buffer. Beads resuspended in the last wash were transferred in a centrifugal device (MilliporeSigma, UFC30LG25) and eluted with the HA peptide at 250 ng/ml two times (30 minutes at 37 °C). In each experiment, three DUBs were tested in parallel with the control condition (no DUB addition). For the initial ten experiments, each DUB was tested in quadruplicate except for the experiment that assessed USP7, USP15 and USP21, in which samples were tested in duplicate. For subsequent follow-up experiments examining USP7, USP36, USP9, USP8 and USP16, each DUB was tested in triplicate with the exception of USP7, which was tested in quadruplicate. Each condition has a volume of extract of 20 μl. Samples were processed for mass spectrometric analysis by SL-TMT method (described below). The following DUBs were tested and purchased from R&D SYSTEMS: E-519 (USP7), E-594 (USP15), E-592 (USP10), E-602 (USP47), E-341(ATXN3), E-572 (YOD1), E-522B (OTUB1), E-621 (MINDY3), E-596 (USP4), E-320 (USP8), E-612 (USP16), E-602 (USP11), E-546 (USP25), E-570 (USP28), E-618 (JOSD1), E-619 (JOSD2), E-624 (OTUD6A), E-322 (USP22), E-522 (USP9X), E-628 (USP36), E-556 (CYLD), E-576 (USP19), E-616 (USP24), E-582 (USP30), E-326 (USP5), E-544 (USP14), E-590 (VCPIP1), E-588 (USP13), E-560 (ZRANB1). Before use, the activity of each DUB was verified with a UbVS-reactivity assay, as previously described^13^. USP13 was the only UbVS-insensitive DUB, as previously reported^23^.

### Streamlined Tandem Mass Tag Protocol

#### Protein extraction and digestion

The TMT labeling protocol and mass spectrometric analysis were based on the SL-TMT sample preparation strategy^51^. Precipitation with trichloroacetic acid (TCA) was performed. 50 μl of TCA were added to 200 μl of the HA-peptide eluate. After incubation of 30 minutes at 4°C, samples were centrifugated for 15 minutes at 4°C (20,000 rcf). Pellets were washed with 1 mL ice-cold acetone, briefly vortexed and centrifugate for 5 minutes at 4°C (20,000 rcf). The acetone wash was then repeated. Next, pellets were washed once with 1mL of cold methanol, vortexed and centrifuged for 5 minutes at 4°C (20,000 rcf). The supernatants were removed by aspiration leaving ∼ 10 μl behind. Pellets were resuspended in 200 mM EPPS, pH 8.5 and digested overnight at 24°C with Lys-C protease (Wako Chemicals, NC9223464). Later, trypsin protease (Pierce Biotechnology, 90058) was added to the samples that were incubated for 6 hours at 37°C. 1 µg of each enzyme was used per 100 µg of protein.

#### TMT Isobaric labeling

10 μL of the tandem mass tag (TMT) isobaric labeling reagents (Thermo Fisher Scientific, 90110) and 30 µL of acetonitrile were added to the digestion mix (∼100 µg). The labeling reaction was incubated at room temperature (90 minutes) and the reaction was then quenched with hydroxylamine to a final concentration of 0.3% (v/v).

#### Quality control analysis of the samples

Prior to proceeding with the final sample pooling and fractionation, we performed a quality control analysis in which a small amount of each sample (2 μl) was pooled together 1:1 and analyzed by mass spectrometry^50^. We confirmed that peptide labeling efficiency was greater than 95%. If all the samples contained the same amounts of peptides, we combined them 1:1 for the final fractionation and analysis. If there were differences among the samples greater than 1.5 fold, we adjusted the relative amounts prior to pooling and fractionation. Following pooling, samples were cleaned up and desalted using a C18-based SepPak cartridge (Waters, WAT054955). The pooled TMT-labeled peptide sample was then fractionated using the Pierce High pH Reversed-Phase Peptide Fractionation Kit (Thermo Fisher Scientific 84868). We followed the manufacturer’s instructions except that the percentage of acetonitrile used in the elution buffers and the subsequent concatenation strategy was performed as described below. Twelve fractions were collected using: 7.5%, 10%, 12.5%, 15%, 17.5%, 20%, 22.5%, 25%, 27.5%, 30%, 35%, and 60% acetonitrile in 0.1% triethylamine (TEA). The samples were then concatenated into six by combining as follows: 7.5% and 22.5%, 10% and 25%, 12.5% and 27.5%, 15% and 30%, 17.5% and 35%, 20% and 60%. Combined fractions were subsequently vacuum centrifuged to near dryness. Each fraction was desalted via StageTip^52^, dried again via vacuum centrifugation, and reconstituted in 5% acetonitrile, 5% formic acid for LC-MS/MS data acquisition.

### Mass spectrometry data collection and analysis

An Orbitrap Lumos or Eclipse mass spectrometer was used to collect the data. The instrument is coupled to a Proxeon NanoLC-1200 UHPLC and in most experiments to a FAIMSpro interface. The sample was analyzed across a 100 μm capillary column packed with 35 cm of Accucore 150 resin (2.6 μm, 150 Å; ThermoFisher Scientific). The spectra from each RAW file were converted to mzXML with MSconvert ^53^ and then searched with the *Xenopus laevis* Database 10.1 plus the same protein sequences with amino acid residues in the reversed order. Our forward databases consisted of 79,357entries. This database was concatenated with a decoy database in which each protein sequence is reversed for the calculation of the false discovery rate using the target-decoy strategy. For real-time search data (RTS), use uses a precursor mass tolerance of 50 ppm and a fragment mass tolerance of 0.9 Da, whereas the fragment mass tolerance is set at 0.03 Da for hrMS2 data. These wide mass tolerance windows were chosen to maximize sensitivity in conjunction with Comet searches and linear discriminant analysis^54^. In addition, oxidation of methionine residues (+15.995 Da) was set as a variable modification, whereas alkylation with N-ethylmaleimide at cysteine residues (+125.048 Da) and TMT or TMTpro tag modifications at peptide N-termini and lysine residues (+229.163 for TMT or +304.207 Da for TMTpro) were set as static modifications. A linear discriminant analysis was performed for PSM filtering, such that a 1% false discovery rate (FDR) for peptide-spectrum matches (PSMs) was set^55,56^, after which then assembled further to a final protein-level FDR of 1% ^57^. The manufacturer-provided isotopic impurities of each TMT or TMTpro reagent were used to correct reporter ion intensities^57^. The peptide signal-to-noise (S/N) measurements for each protein were summed.

### Confirmation of candidate DUB substrates with S35-labeled substrates

Substrates were expressed and labeled using ^35^S-methionine (Perkin Elmer, NEG709A500UC) with the T7 TNT Coupled Reticulocyte Lysate System (Promega, L1170). Each substrate was amplified with primers by PCR (using template plasmids from the hORFeome collection or commercially available plasmids) to allow T7-dependent transcription of the PCR product. The translation reaction mix was added to *Xenopus* extract at 8% final volume in the presence of UbVS (10 μM). After 30 min at 24°C, 50 μM of human recombinant untagged ubiquitin (R&D SYSTEMS, U-100H) and specific human recombinant DUBs were added at the same concentration as for the proteomic experiments (800 nM). Samples of the reactions were collected at the indicated time, quenched with SDS sample buffer, and processed for SDS gel electrophoresis and phosphor imaging. Images were acquired with the Typhoon laser-scanner platform (Cytiva) and quantified with ImageQuantTL (Cytiva).

## QUANTIFICATION AND STATISTICAL ANALYSIS

For the TMT-based mass spectrometric analysis, a two-sided Student’s t-test was used as a measure of statistical confidence of the observed log_2_ fold change. Selected candidates met both thresholds (fold Change < −0.5 and *p*-value<0.05). For statistical analysis, t-test or Chi squared test were performed as indicated in the figure legends.

### Creation of an interactive web-based application to easily access and explore our data

To make our data easily available, we developed a R-shiny application which allows users to independently explore all the data we presented in our study.

### Our application offers different options to query the data

In the *single protein profile* tab, the user can search for a protein of interest to check the effect of each DUB on the selected protein. As an example, the profile for the ubiquitin ligase TRIP12, a known substrate of USP7^10^ that we also identified as a candidate substrate for USP7 is shown in Fig. 8A and Fig. 8B.

On the *volcano plot* tab, the user can visualize the volcano plot or the list of candidate substrates identified for that DUB. The user simply types the name of the DUB of interest and the experiment number in which the DUB was tested to visualize the volcano plot or the table of the candidate substrates identified for that DUB. The volcano plot compares the immunopurified proteins detected on the beads after addition of the DUB of interest with the immunopurified proteins detected in absence of any DUB (log_2_ DUB/NO DUB). The USP7 example is shown in Fig 8C. Near the volcano plot, a table reports all the proteins of the volcano plot and their values.

On the *list of candidate substrates* tab, the user can explore the list of candidate substrates for a DUB of interest with links to the String, Uniprot, and D2P2 database (disorder), to retrieve information about each substrate (Fig. 8D).

On the *hierarchical clustering* tab, the user can globally explore our data in the form of a two-way hierarchical cluster analysis based on the activity of each DUB against the common set of proteins (Fig. 8F) or against the entire dataset.

